# SARS-CoV-2 Vaccines Elicit Durable Immune Responses in Infant Rhesus Macaques

**DOI:** 10.1101/2021.04.05.438479

**Authors:** Carolina Garrido, Alan D. Curtis, Maria Dennis, Sachi H. Pathak, Hongmei Gao, David Montefiori, Mark Tomai, Christopher B. Fox, Pamela A. Kozlowski, Trevor Scobey, Jennifer E. Munt, Michael L. Mallroy, Pooja T. Saha, Michael G. Hudgens, Lisa C. Lindesmith, Ralph S. Baric, Olubukola M. Abiona, Barney Graham, Kizzmekia S. Corbett, Darin Edwards, Andrea Carfi, Genevieve Fouda, Koen K. A. Van Rompay, Kristina De Paris, Sallie R. Permar

**Author notes:** **equal first authors** (Carolina Garrido, Alan D. Curtis). **equal senior authors** (Kristina De Paris, Sallie R. Permar).

## Abstract

Early life SARS-CoV-2 vaccination has the potential to provide lifelong protection and achieve herd immunity. To evaluate SARS-CoV-2 infant vaccination, we immunized two groups of 8 infant rhesus macaques (RMs) at weeks 0 and 4 with stabilized prefusion SARS-CoV-2 S-2P spike (S) protein, either encoded by mRNA encapsulated in lipid nanoparticles (mRNA-LNP) or mixed with 3M-052-SE, a TLR7/8 agonist in a squalene emulsion (Protein+3M-052-SE). Neither vaccine induced adverse effects. High magnitude S-binding IgG and neutralizing infectious dose 50 (ID_50_) >10^3^ were elicited by both vaccines. S-specific T cell responses were dominated by IL-17, IFN-*γ*, or TNF-*α*. Antibody and cellular responses were stable through week 22. The S-2P mRNA-LNP and Protein-3M-052-SE vaccines are promising pediatric SARS-CoV-2 vaccine candidates to achieve durable protective immunity.

**One-Sentence Summary:** SARS-CoV-2 vaccines are well-tolerated and highly immunogenic in infant rhesus macaques

## Main text

Severe acute respiratory syndrome coronavirus 2 (SARS-CoV-2) has infected over 110 million people worldwide and caused close to 2.5 million deaths since its emergence in 2019. The need for safe and effective measures to limit transmission and mitigate public health and socioeconomic impacts of SARS-CoV-2 infection has prompted unprecedented vaccine development of promising candidates.

Two messenger RNA (mRNA) vaccines, mRNA-1273 (Moderna) (*1, 2*) and BNT162b2 (Pfizer-BioNTech) (*3, 4*), have been authorized for emergency use in the United States to prevent SARS-CoV-2 infection. Both demonstrated safety, strong induction of neutralizing antibodies and up to 95% protection from disease. Virus-vectored vaccine platforms, including the Oxford/AstraZeneca adenovirus-based ChAdOx1 nCoV-19 (AZD1222) (*5*) and the Johnson&Johnson Ad26.COV2.S vaccine (JNJ-78436735) (*6*), and the first protein-based SARS-CoV-2 vaccine, Novavax NVX-CoV2373, adjuvanted with saponin-based Matrix-M™ (*7, 8*), have also demonstrated high efficacy in clinical trials.

Early in the pandemic, vaccines to prevent of SARS-CoV-2 infection in children were not a priority because of apparent low infection and disease rates. However, we now know that children can contract SARS-CoV-2 as frequently as other age groups (*9*). Despite reduced hospitalizations and deaths in children compared to adults, some children, especially those with comorbidities, develop severe disease, including multisystem inflammatory syndrome (*10, 11*). The important epidemiologic impact of pediatric SARS-CoV-2 vaccination lies in limiting transmission and ease of implementation via the global pediatric vaccine schedule. Yet, to date only children 12 years and older have been recruited in a few trials (NCT04649151, NCT04368728) with a single trial in early enrollment for ages 6 months to 12 years (NCT04796896). However, vaccine-elicited immunity differs between adults and infants (reviewed in (*12*)), with impaired responses to carbohydrate antigen vaccines (*13*), but higher magnitude humoral immunity to subunit-based vaccines (*14*) early in life. Thus, the evaluation of SARS-CoV-2 vaccine immunogenicity in infants is critical.

Non-human primates (NHP) are an important model for SARS-CoV-2 studies given the similarities in pathogenesis and host immune responses (*15*) to humans. Indeed, results from adult NHP studies of human SARS-CoV-2 vaccine candidates (*16–19*) strongly correlate with clinical trial outcomes, and infant NHP models of other infectious diseases closely mirror responses in human infants (*20, 21*). The present study was designed to evaluate the safety and immunogenicity of stabilized prefusion SARS-CoV-2 Spike vaccines delivered as lipid nanoparticle-encapsulated mRNA or an adjuvanted subunit protein in infant rhesus macaques (RMs).

### Study design and vaccine safety

Infant RMs at a median age of 2.2 months, corresponding to 9-month old human infants (*22*), were immunized at weeks 0 and 4 with 30 µg mRNA encoding stabilized prefusion SARS-CoV-2 S-2P protein in lipid nanoparticles (mRNA-LNP; n=8) or with 15 µg S-2P mixed with 3M-052-SE, a TLR7/8 agonist in a stable emulsion (Protein+3M-052-SE; n=8) (Fig. 1, Table S1). Animals were monitored daily for adverse events and did not experience injection site swelling, allergic reactions, had normal blood metrics (Table S2), and weight gain consistent with normal infant growth (Fig. S1). Blood and saliva were collected before vaccination (W0), 4 weeks after the first dose of the vaccine (W4), 2 weeks after the vaccine boost (W6), and at weeks 8, 14, 18 and 22. Lymph nodes were sampled at week 6 (Fig. 1).

**Figure 1.**
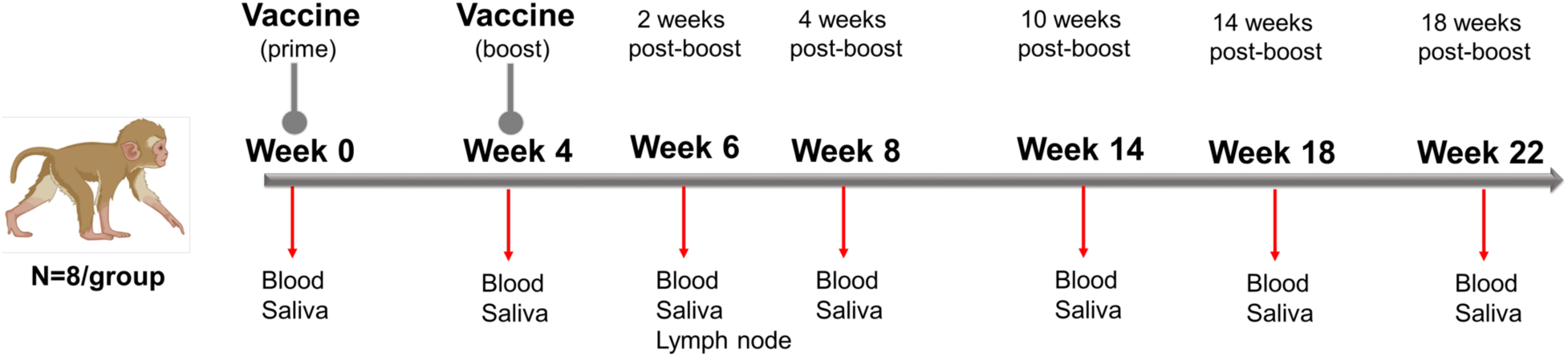
Study Design: evaluation of immunogenicity of two SARS-CoV-2 vaccines in infant rhesus macaques. Infant rhesus macaques (median age of 2.2 months at study initiation) were immunized at 0 and 4 weeks with either 30 µg mRNA encoding S-2P (Vaccine Research Center, NIH) in lipid nanoparticles (mRNA-LNP) or 15 µg S-2P protein formulated with 3M-052 adjuvant, a TLR7/8 agonist, as a stable emulsion (3M-052-SE). Each group consisted of 8 animals. Blood and saliva samples were collected at weeks 0, 4, 6, 8, 14, 18 and 22, and lymph node biopsies were obtained at week 6.

### Infant antibody responses to SARS-CoV-2 vaccination

Plasma IgG binding to the S-2P protein was observed after the vaccine prime for both mRNA-LNP and Protein+3M-052-SE vaccines and increased after the second immunization. Although antibody levels dropped by week 8, there was only a slight decline of S-2P-specific IgG from week 8 to week 22 for mRNA-LNP (median AUC and 95% confidence intervals: 7.0 [6.2, 8.3] to 5.4 [4.2, 7.7]) or Protein+3M-052-SE (11.0 [9.5, 11.7] to 9.4 [8.6, 10.5] vaccinees (Fig. 2A). As the D614G virus variant had become dominant by study initiation, we confirmed that IgG binding to S-2P D614G was similar to that of D614 at weeks 6 and 14 (Fig. S2). Plasma IgG were directed against multiple spike domains, with robust binding to Spike regions 1 and 2 (S1, S2), Receptor binding domain (RBD), and N-terminal domain (NTD), that persisted throughout the study (Fig. 2B). RBD-specific salivary IgG from mRNA-LNP recipients peaked at a median of 16.6 ng RBD-specific IgG per µg of total IgG at week 6 (Fig. 2C). In the Protein+3M-052-SE group, median salivary RBD-specific IgG peaked at 98.2 ng/µg IgG after the second vaccination and remained detectable throughout the study (Fig.2B). Saliva RBD-specific IgA responses were much lower (Fig S3). Spike-specific IgM and IgA in plasma were undetectable or low (Fig. S3). In the mRNA-LNP group, IgM increased through week 14, but the absorbance for IgM are considered low, measured at a 1:10 sample dilution (Fig. S3).

**Figure 2:**
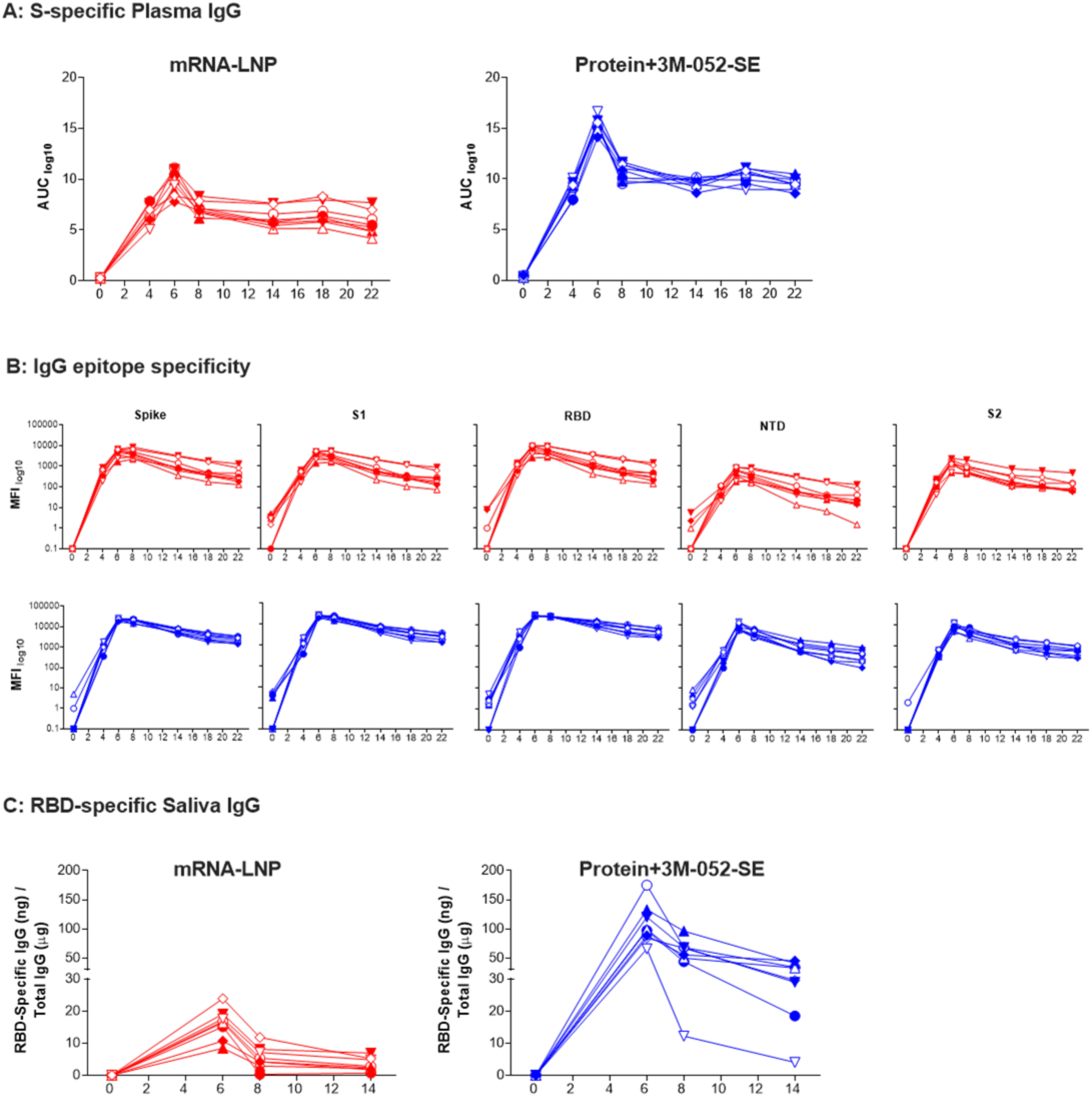
SARS-CoV-2 vaccine-elicited binding antibody responses in infant rhesus macaques. Plasma and saliva were collected before vaccination (W0), at W4 -just prior to the boost-, two weeks post boost (W6), at W8, W14, W18 and W22 from infant RM vaccinated with 30 µg mRNA encoding S-2P spike protein in lipid nanoparticles (n=8; red) or with 15 µg prefusion SARS-CoV-2 S-2P spike protein formulated with 3M-052 adjuvant (n=8; blue). **(A):** S-2P protein-specific antibody responses were measured by enzyme-linked immunosorbent assay (ELISA). Serial dilutions of plasma starting at 1:40 were assayed for IgG binding to SARS-CoV-2 spike. Data are reported as log_10_ area under the curve (AUC) values. **(B):** Salivary RBD-specific IgG was measured by binding antigen multiplex assay (BAMA) using serial dilutions of saliva. **(C):** Antibody epitope specificity measured by BAMA. Plasma was diluted 1:10,000 to measure binding to different domains of the spike protein, including the full-length S protein, S1, RBD, NTD, and S2. Binding antibody responses are reported as log_10_ transformed mean fluorescence intensity (MFI) after subtraction of background values. Red or blue lines and symbols represent the mRNA or protein vaccine groups, respectively, with different symbols representing individual animals (Table S1).

We evaluated the capacity of plasma antibodies to block entry of SARS-CoV-2 into human cells using a RBD-ACE2 blocking assay at 1:10 and 1:40 plasma dilutions. Antibodies elicited by the mRNA-LNP vaccine completely blocked RBD-ACE2 interaction at week 6 at 1:10 and 1:40 dilutions. Until W14, >80% blocking was achieved at 1:10 dilution, dropping for some animals after W18 (Fig. 3A). The Protein+3M-052-SE vaccine induced antibodies that mediated 100% RBD-ACE2 blocking after the second immunization, even at 1:40 plasma dilution, and this response was maintained throughout the study in 5 of 8 animals (Fig. 3A, Fig. S4). Soluble ACE2 protein, as measured by ELISA, was absent from infant plasma through week 22 (data not shown).

**Figure 3:**
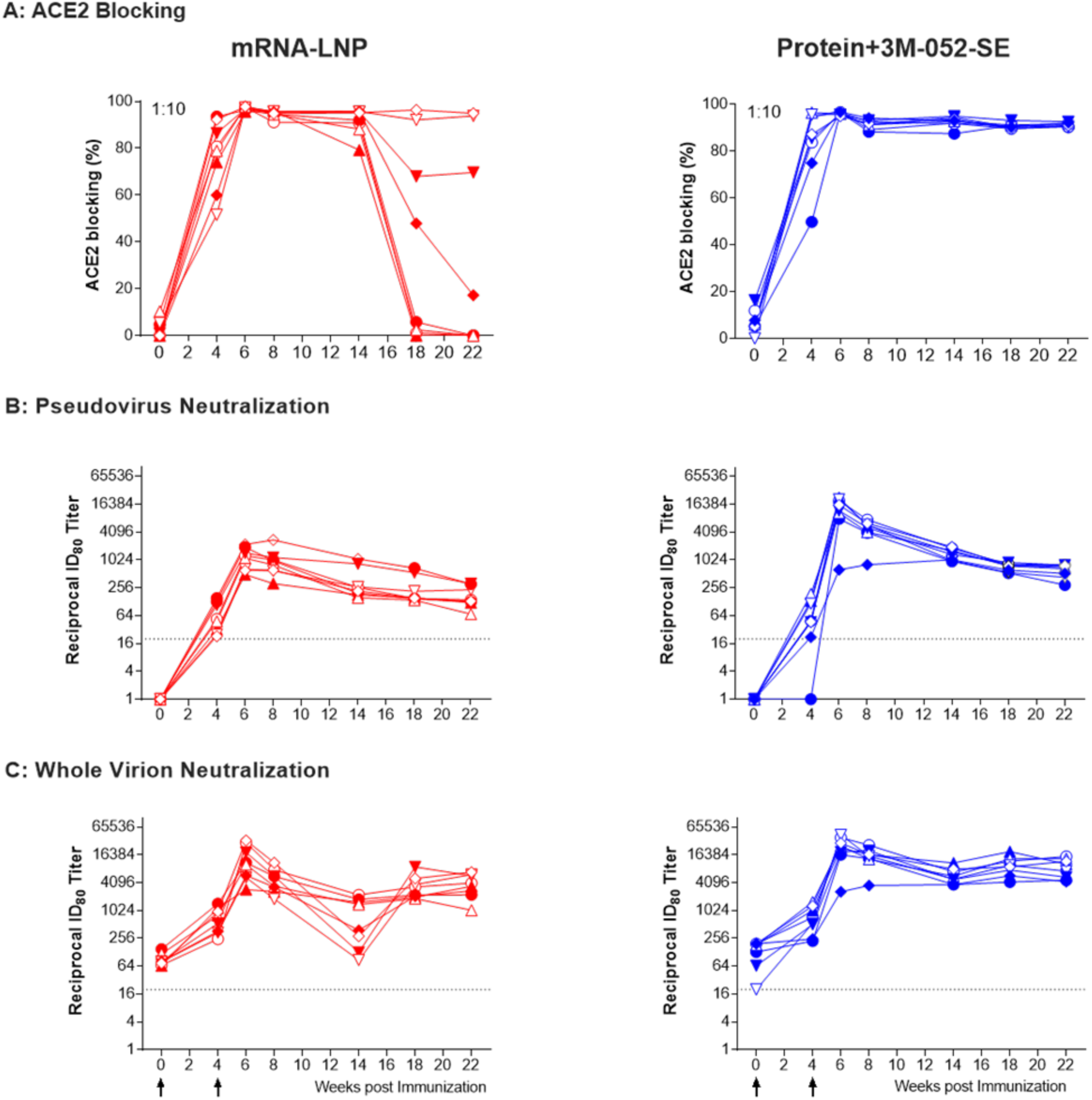
SARS-CoV-2 vaccine-elicited functional antibody responses in infant rhesus macaques. **(A):** The ACE2 blocking assay was performed at 1:10 plasma dilution and data are reported as %ACE2 blocking. **(B-C)**: Neutralization capacity was measured using a pseudovirus assay **(B)** and whole virus assay **(C)**; results are expressed as reciprocal 80% inhibitory dilution (ID_80_). Different symbols represent individual animals (Table S1).

Neutralizing antibodies, measured by a pseudovirus assay, were detected in animals of both groups after the first vaccination, and further increased after the boost. At week 6, median infectious dose 80 (ID_80_) titers, a stringent neutralization measurement, reached 1,179 in mRNA-LNP and 13,687 in Protein+3M-052-SE animals (Fig. 3B). Consistent with normal contraction of immune responses, median ID_80_ neutralization titers decreased 1.3-fold and 2.6-fold from week 6 to week 8 in the mRNA-LNP and Protein+3M-052-SE group, respectively. However, at week 22, median ID_80_ titers remained higher (2.6-fold or 14.2-fold in the mRNA-LNP or Protein+3M-052-SE group, respectively) than after the first vaccination (week 4) (Fig. 3B). A bi-phasic antibody decline was modeled for both vaccines, estimating that mRNA-LNP vaccinees maintain a protective neutralizing Ab titer, estimated at a reciprocal dilution of 100, for approximately 46 weeks. Protein vaccine recipients sustain protective levels for approximately 74 weeks (Fig. S5). Importantly, ID_50_ neutralization titers in the mRNA-LNP group at week 8 were comparable to those observed in adult RMs after immunization with the same S-2P mRNA-LNP vaccine (*16*) (Fig. S6). In the whole virus neutralization assay, low titer neutralizing antibodies were detected at day 0 that increased post vaccination following a similar trend (Fig.3C), and ID_80_ titers correlated with those obtained in the pseudovirus neutralization assay (r = 0.785, p=0.0005, Fig. S6). Whole virion neutralization was also measured in dam sera, observing some low-level neutralization titers that did not correlate to infant neutralization before (ID_80_: r=0.011, p=0.9681) or after vaccination (ID_80_: r=-0.056, p=0.8362) (Fig. S7).

### Vaccine-elicited B and T cell responses

S-specific memory CD27^+^ B cell were detectable in blood of both vaccine groups at week 4 and peaked at week 6 (median: 075% mRNA-LNP; 3.12%, Protein+3M-052-SE). At week 14, median memory B cell frequencies were lower (0.19% and 0.13% in mRNA-LNP or Protein+3M-052-SE vaccinees, respectively), but did not differ from those at week 4 (Fig. 4A). Robust memory B cell populations (median 3.78% mRNA-LNP and 1.96% Protein-3M-052) were present in lymph nodes (LNs) (Fig. 4B). Induction of S-specific B cells was confirmed by assessing S-specific antibody secreting cells (ASC). In blood, peak responses were observed at week 6 in the mRNA-LNP group (median: 154 ASC/million cells) and at week 8 in Protein+3M-052-SE vaccinees (median: 85 ASC/10^6^ cells); S-specific ASC remained detectable at all time points in both groups (Fig. 4C). In draining LNs, mRNA-LNP and Protein+3M-052-SE vaccinees had median 6 or 952 ASC/10^6^ cells, respectively (Fig. 4D). Combined data from both vaccine groups showed neutralizing ID_50_ titers at weeks 6 and 8 correlated with ASC (r=0.547, p=0.03 and r=0.632, p=0.01, respectively) (Fig. S9A-B), but not with LN S-specific memory B cells. Total LN Ki-67^+^Bcl6^+^ germinal center (GC) and CD27^+^ memory B cells (Fig. 5A) were also not correlated with neutralizing antibody titers.

**Figure 4:**
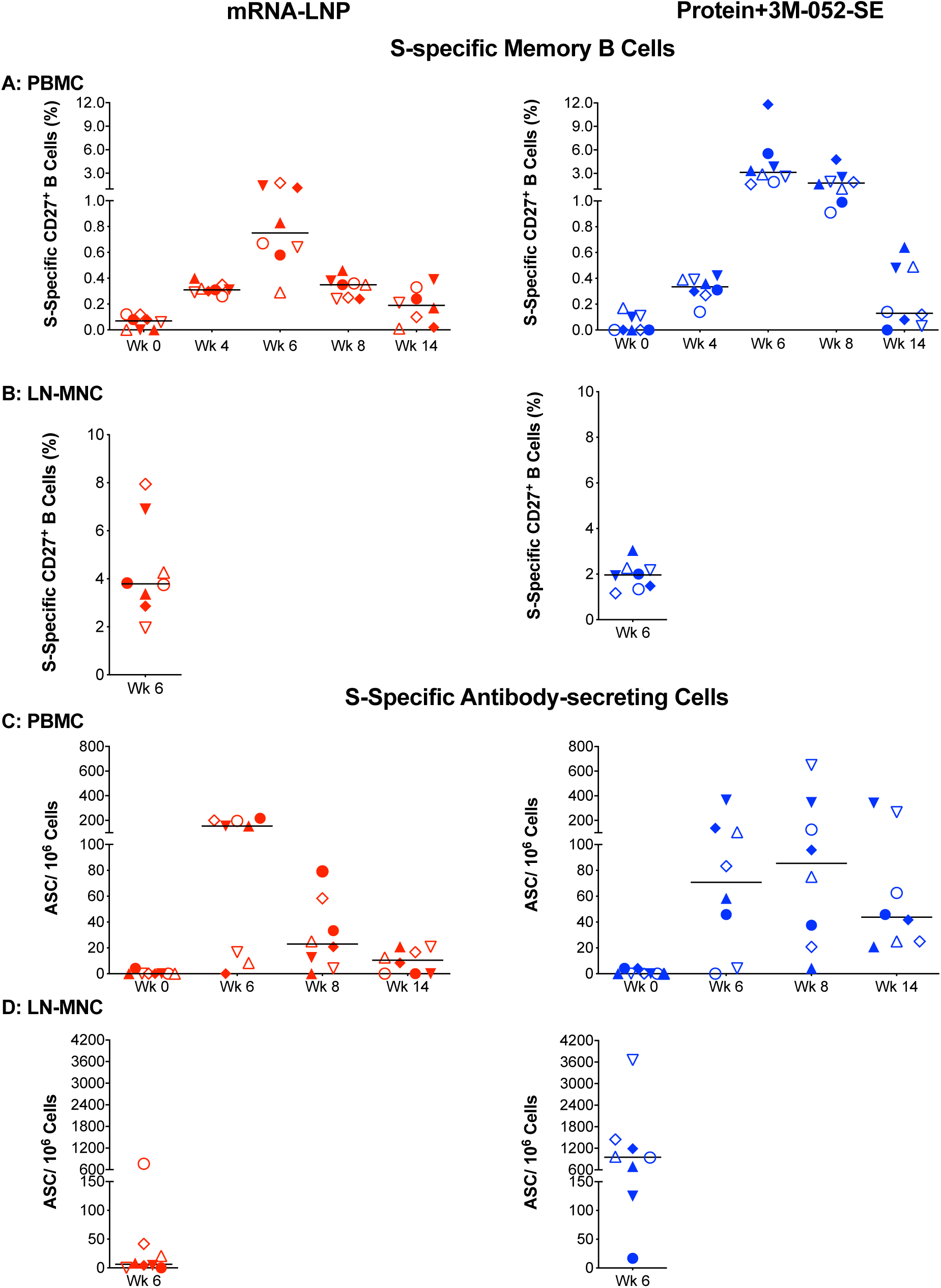
Characterization of Spike-specific B cell responses two weeks post boost. CD20^+^CD27^+^ memory B cells that co-stained with fluorochrome-conjugated SARS-CoV-2 spike protein in mRNA-LNP (red) or Protein+3M-052-SE (blue) vaccinees in blood (**A - B**) or LN (**C - D**). Frequencies are expressed as percent of total memory B cells. The gating strategy is provided in Supplementary Figure S8. (**E–F)** portray antibody secreting cell (ASC) as measured by B cell ELISpot in PBMC from mRNA-LNP or Protein+3M-052-SE vaccinees, while **(G–H)** contain mRNA-LNP and Protein+3M-052-SE ASC responses, respectively, in LN at W6. Different symbols represent individual animals (Table S1). Solid lines represent median values.

**Figure 5.**
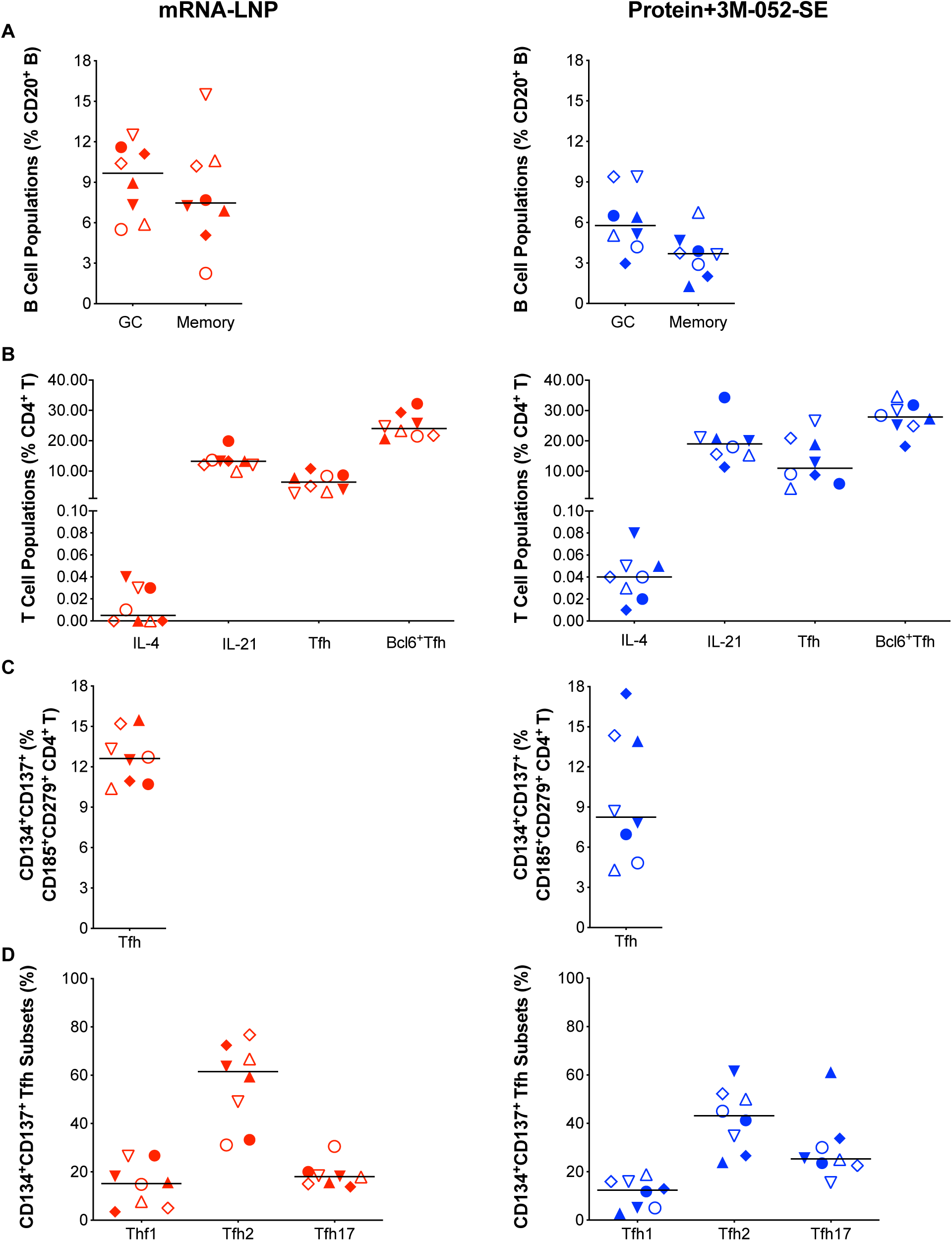
Immunophenotype of lymph node T and B cell population two weeks post-boost. **(A–B):** Frequencies of Bcl-6^+^Ki-67^+^ Germinal Center (GC) B cell and CD27^+^ memory B cells as percent of total CD20^+^ B cells from mRNA-LNP or Protein+3M-052-SE recipients, respectively. **(C–D):** CD4^+^ T-cells positive for IL-4 or IL-21, follicular T helper cell (Tfh) markers CD279/PD-1 and CD185/CXCR5, or Bcl6^+^ Tfh were measured and are represented as percent of total LN CD4^+^ T cells. The gating strategy for these panels is provided in Figure S10. **(E–F)** show SEB activated Tfh frequencies assessed by the Activation-Induced Marker (AIM) Assay: CXCR5^+^CD185^+^ cells that co-expressed CD134 and CD137 for Protein+3M-052-SE and mRNA-LNP, respectively. **(G–H)** depict the frequency of CXCR5^+^CD185^+^CD134^+^CD137^+^CXCR3^+^CD196^-^ Tfh1, CXCR3^-^CCR6^-^ Tfh2 and CXCR3^-^CD196^+^ Tfh17 cells. The gating strategy for these populations is given in Figure S3. Red symbols: mRNA-LNP, blue symbols: Protein+3M-052-SE. Different symbols represent individual animals (Table S1). Solid lines define the median.

LN CD4^+^ T cells were analyzed for IL-4 and IL-21, cytokines supporting GC B cell differentiation, and canonical T follicular helper (Tfh) (CD185/CXCR5^+^CD279/PD-1^+^) cells (Fig. 5B). CD4^+^IL21^+^ cells correlated with pseudovirus neutralizing antibody ID_50_ at week 6 (r=0.749, p=0.001) (Fig. S9C). SEB-activated Tfh (CD185^+^CD279^+^CD134^+^CD137^+^) (*23*) (Fig. 5C) were categorized into Tfh1, Tfh2 and Tfh17 subsets based on expression of CXCR3/CD183 and/or CCR6/CD196 (Fig. 5D). We found no correlations between frequencies of Tfh subsets with neutralizing antibody titers, GC or memory B cells, potentially because we measured antigen-nonspecific Tfh.

S-specific CD4^+^ T cell responses in PBMC could already be detected by week 4 in a subset of animals, with all animals producing at least a single cytokine by week 6 (Fig. 6). At week 14, animals vaccinated with mRNA-LNP exhibited IL-2, IFN-*γ*, IL-17, and TNF-*α* CD4^+^ T cell responses (Fig. 6D). IL-17 and IFN-*γ* CD4^+^ T cell responses dominated in Protein+3M-052-SE vaccinees (Fig. 6D). Multifunctional CD4^+^ T cells co-produced IL-17 and IFN-*γ* (Fig. S11). S-specific CD8^+^ T cell responses were less robust than CD4^+^ T cell responses and only elicited in a subset of animals (Fig. S12). LN S-specific CD4^+^ and CD8^+^ T cells were single-cytokine positive and detected in all mRNA-LNP recipients at week 6 and in 7 of 8 Protein+3M-052-SE vaccinees (Fig. S13). There was no evidence of systemic IL-2, IFN-γ, IL-4, or IL-13 or in plasma samples collected prior to or following immunization in 14 of 16 animals (Fig. S14). Cytokines were detected in the mRNA-LNP recipient RM11 (Table S1) at week 0, but were below the limit of quantification by week 6. In the protein group, animal RM3 (Table S1) tested positive for all cytokines at both time points. Overall, the data argue against a vaccine-induced Th2 bias.

**Figure 6:**
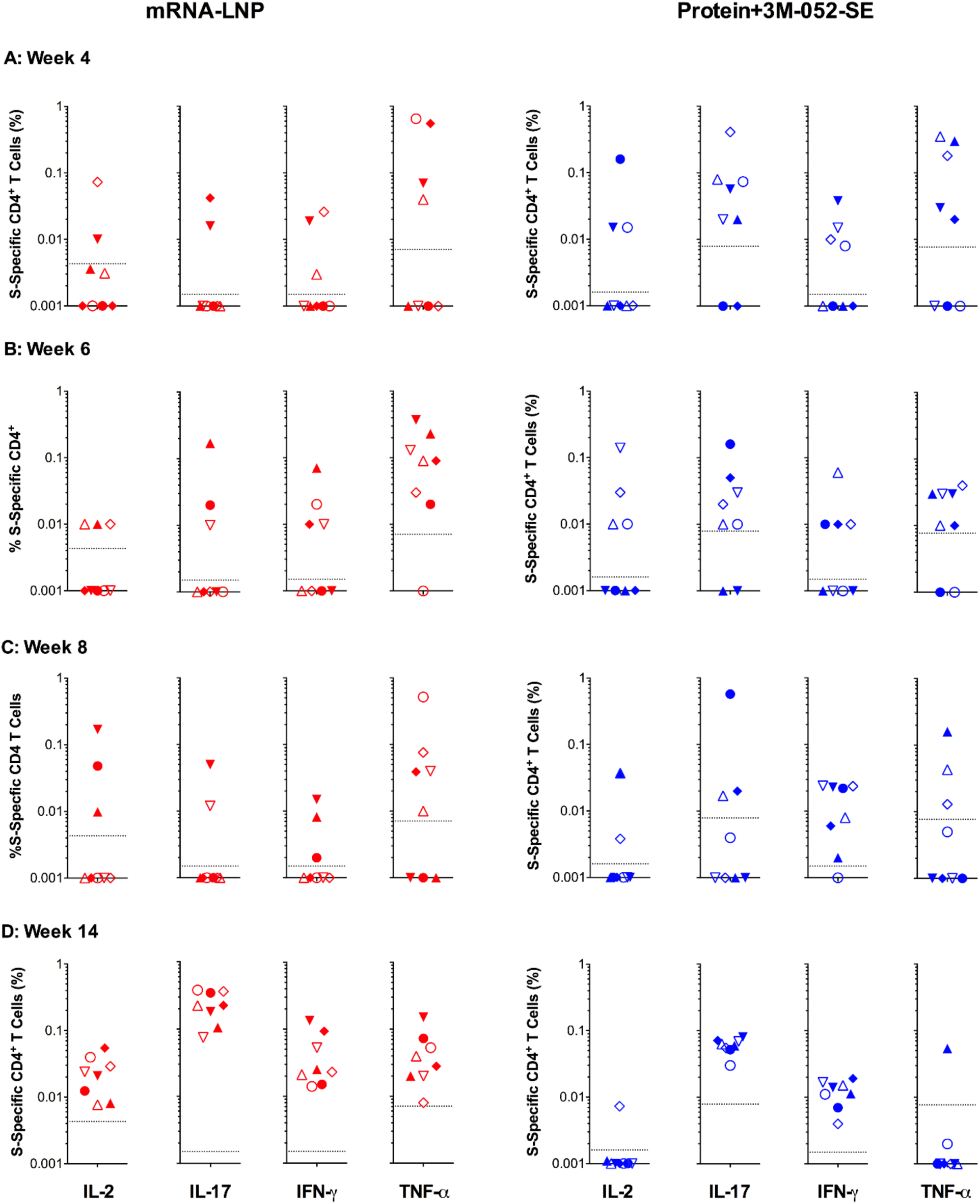
Spike-specific CD4^+^ T cell responses in SARS-CoV-2 immunized infant macaques. Intracellular cytokine staining for IL-2, IL-17, IFN-γ, and TNF-α was performed on PBMC at weeks 0, 6, 4, 8, and 14 to assess T-cell responses to a peptide pool encompassing the entire SARS-CoV-2 spike protein. **(A–D)** display responses detected in Protein+3M-052+SE vaccinees (blue). **(E-H)** portray cytokine responses from mRNA-LNP recipients (red). The dashed lines represent week 0 values plus 2 standard deviations and define the cutoff for positive cytokine responses. Different symbols represent individual animals (Table S1).

## Discussion

Several SARS-CoV-2 vaccines have demonstrated safety, immunogenicity and protection in clinical trials (*1, 5, 6*) and adult animal studies (*8, 16, 24, 25*). However, to date there are no data regarding the performance of these vaccines in pediatric populations. We evaluated two vaccines based on the stabilized prefusion S-2P SARS-CoV-2 protein in infant RMs: a LNP-encapsulated mRNA vaccine and an adjuvanted subunit protein. Our results demonstrate that infant RMs develop strong, durable humoral and cellular responses in the absence of adverse events following vaccination with either vaccine candidate.

We assessed infant vaccine-induced immune responses over 22 weeks, analogous to adult clinical trials (*1, 2*). The S-2P mRNA-LNP vaccine, a preclinical version of the highly efficacious mRNA-1273 vaccine by Moderna, that is approved for emergency use in adult humans, elicited high magnitude and durable antibody and cellular responses in infant RMs. Efficacy of the first protein-based SARS-CoV-2 vaccine, Novavax NVX-CoV2373, mixed with saponin-based Matrix-M™ adjuvant, has recently been reported (*8*). We tested the S-2P SARS-CoV-2 protein mixed with 3M-052-SE, a TLR7/8 agonist in squalene emulsion, because this adjuvant has proven effective in eliciting high magnitude antibody responses to other vaccines in infant RMs (*26, 27*). Indeed, the 3M-052-SE formulated S-2P protein induced robust and persistent IgG binding and neutralization responses.

Both vaccines elicited plasma antibodies dominated by IgG and recognized all Spike protein domains. Interestingly, RBD-specific IgG was also detected in saliva, especially in animals of the protein vaccine group, and remained detectable throughout the study, similar to what has been observed after human natural infection (*28*). Moreover, 7 of 8 Protein-3M-052-SE vaccinated animals had RBD-specific IgA after the 2^nd^ immunization. Salivary RBD-specific IgG and IgA may either be mucosally produced or transudated from plasma.

Binding and neutralizing antibodies persisted throughout the study in all animals of both groups, comparable to results in clinical trials with human adults. In adult NHP, a vaccine-induced pseudovirus neutralizing antibody ID_50_ titer of 50 is thought to protect against susceptible SARS-CoV-2 strains (*29*). As median ID_50_ titers exceeded 10^3^ for the protein vaccine group and 10^2^ for the mRNA group at week 22, we hypothesize that pediatric vaccination would provide long-term protection against SARS-CoV-2 infection, with the potential need to include additional S variant antigens of newly emerging SARS-CoV-2 variants into vaccine candidates as necessary.

Early life immunity is associated with Th2-biased T cell responses (*30*), and Th2 responses have been linked to vaccine-associated enhanced respiratory disease in the context of protein or inactivated virus vaccines (*6, 31, 32*). We observed a Th1/ Th17-skewed cytokine profile in circulating S-specific T cells. Our finding of peak S-specific T cell responses 10 weeks post-boost are not unexpected. A recent study of T cell reactivity in convalescent COVID-19 patients highlighted T cell responses with augmented frequency and potency 100 days post-recovery even as S-specific antibody waned (*33*). Indeed, ∼93% of “exposed asymptomatic” patients possess SARS-CoV-2-specific memory T cell subsets in the absence of seroconversion (*34*). Higher Tfh2 frequencies compared to Tfh1 does not necessarily imply a Th2-bias as Tfh17 cells were also detected, and Tfh were not assessed for antigen specificity. More importantly, IL-4 and IL-21 producing Tfh are required for the induction of GC B cells in LNs (*35*), indispensable for B cell and antibody affinity maturation, and GC reactions are important for positive outcomes in COVID-19 patients (*36*). Indeed, the persistence of antibody was paralleled by sustained S-specific B cell responses. The induction and persistence of S-specific B cell clones described here may be key to protection against re-infection (*37*). We found no evidence of systemic Th2 cytokines prior to or following immunization in either vaccine group, corroborating previous findings in adult macaques (*16*) and adult humans (*38*) that also demonstrated low level or absent Th2-mediated responses.

This is the first report of pediatric immune responses to SARS-CoV-2 vaccination. Our results demonstrate that infant RMs mount strong and durable responses to mRNA-LNP and protein-based SARS-CoV-2 vaccines that are comparable to adults without adverse reactions, justifying urgent clinical translation of SARS-CoV-2 vaccines to early life populations.

## Acknowledgements

We thank Jennifer Watanabe, Jodie Usachenko, Ramya Immareddy, the Veterinary staff and Colony Research Services of the California National Primate Research Center for expert assistance.

We thank Elise Larson, Ines De Lima, Tony Phan, Susan Lin, and Robert Kinsey from IDRI for technical assistance in preparing and characterizing 3M-052-SE.

We thank Jim Giron of ThermoFisher Scientific for expert technical assistance with T cell cytokine Luminex assays.

We thank Dr. James Peacock in the Duke Human Vaccine Institute Research Protein Production Facility.

## Funding

National Institutes of Health grant P01 AI117915-06S1 (SRP, KDP)

National Institutes of Health grant U54 CA260543 (RB)

National Institutes of Health Office of Research Infrastructure Program/OD P510D11107 (CNPRC)

National Institutes of Health grant UM1 AI068618-15 (HVTN/HPTN, CoVPN)

National Institutes of Health grant P30AI050410 (UNC Center for AIDS Research)

P30 CA016086 (UNC-LCCC Flow Cytometry Core Facility)

Dr. Corbett is the recipient of a research fellowship that was partially funded by the Undergraduate Scholarship Program, Office of Intramural Training and Education, Office of the Director, NIH.

## Author Contributions

Writing -original draft: CG, ADC, SHP, KDP

Writing –review and editing: CG, ADC, MT, CBF, PAK, RSB, BG, KSC, DE, AC, GF, KKAVR, KDP, SRP

Methodology: CG, ADC, MD, SHP, GF, PAK, HG, DM, TS, JEM, MLM, LSL, RSB

Statistical data analysis: PTS, MGH

Animal studies: KKAVR. OB, KC

Vaccine supply: BG, DE, AC, MT, CBF. All authors assisted in editing of the manuscript.

## Supplementary Materials

### Materials and Methods

#### Animals

Infant male (n=8) and female (n=8) rhesus macaques (*Macaca mulatta;* RM) of Indian-origin from the California National Primate Research Center (CNPRC, Davis, CA) breeding colony (negative for type D retrovirus, simian immunodeficiency virus, simian lymphocyte tropic virus type 1 and SARS-CoV-2), were enrolled at a median age of 2.2 months and randomly assigned into two groups (Table S1). Infants were housed with dams in indoor housing in accordance with the “Guide for Care and Use of Laboratory Animals” by the Institute for Laboratory Animal Research. Animal procedures were approved by the UC Davis Institutional Animal Care and Use Committee prior to study initiation. All procedures were performed under anesthesia (ketamine, 10 mg/kg body weight, intramuscularly [IM]). Blood, serum, saliva and lymph nodes (LN) were collected and processed as described (*39*).

#### Vaccines

The SARS-CoV-2 stabilized prefusion Spike (S-2P) mRNA-LNP vaccine was provided by Moderna, Inc. and the Vaccine Research Center (NIH) provided the S-2P protein (see Appendix). The vaccine regimen and specimen collection are outlined in Figure 1. Animals in the mRNA-LNP vaccine group were immunized IM with 30 µg mRNA encoding S-P2 protein in lipid nanoparticles (mRNA-LNP), that was diluted in 0.1 mL phosphate-buffered saline to get to single injection site concentration, at weeks 0 (quadriceps) and 4 (biceps). Infant RMs in the protein vaccine group were injected IM with15 µg S-2P protein mixed with 3M-052-SE, an adjuvant formulation consisting of 10 µg of the synthetic TLR7/8 agonist 3M-052 in a 2% v/v squalene-in-water emulsion (Protein+3M-052-SE) in 0.5 mL divided across the left and right quadriceps (week 0) or biceps (week 4).

#### Antibody measurements

IgG binding to S-2P (2019-nCoV) and the D614G variant was measured in plasma using enzyme-linked immunosorbent assay (ELISA), following published protocols (*40*). IgM and IgA binding to S-2P was measured in 1:10 diluted plasma by ELISA. To assess epitope-specificity, IgG binding to the RBD, NTD, S1, S2 and the whole spike, was measured by a Luminex-based binding antibody multiplex assay (BAMA) at a dilution of 1:10,000 (*40*). RBD-specific IgG and IgA in saliva were quantified in serially diluted samples using BAMA, and results normalized to total IgG and IgA (see Supplementary Material for more details).

RBD-ACE2 blocking by plasma antibodies was measured in 384-well plates coated with human ACE2 protein. Diluted plasma samples (1:10, 1:40 and 1:60) were pre-incubated with horseradish peroxidase (HRP)-conjugated RBD antigen and added to the coated plate. Binding was detected by colorimetric signal after addition of the chromogenic substrate. Wells without plasma were considered maximal binding, and absorbances were normalized to that value and expressed as percentage of binding inhibition.

Pseudovirions were produced in HEK 293T/17 cells by transfection using a combination three plasmids: full-length Spike with D614G mutation, lentiviral backbone (pCMV ΔR8.2) and firefly Luc reporter gene (pHR’ CMV Luc) (*41*). Luminescence was measured using a PerkinElmer Life Sciences, Model Victor2 luminometer after addition of Bright-Glo luciferase reagent. Neutralization activity is expressed as the reciprocal dilution at which the relative light units (RLU) were reduced by 50% (ID_50_) or 80% (ID_80_) relative to virus-only control wells. Whole virus neutralization was performed with a SARS-CoV-2 nanoLUC carrying the D614G mutation as described (*42*).

#### Soluble ACE2 and cytokine measurement

Plasma was evaluated for soluble ACE2 protein with a human plasma ACE2 ELISA kit (Raybiotech) following the manufacture’s guidelines. IL-2, IL-4, IL-13, and IFN-γ in undiluted plasma samples were quantified via custom 4-plex rhesus macaque Luminex assay (Thermofisher Scientific) using protocols established by the supplier. ELISA results are reported as the average concentration of duplicate wells extrapolated from a standard curve.

#### B Cell Responses

S-specific B cells from PBMC and LN were assessed by flow cytometry as described (*27*) (Table S3). Total CD20^+^ and/or memory CD27^+^ S-specific B cells were identified as double-positive for biotinylated S-2P protein labeled with avidin-APC or avidin-PE (Fig. S8). Germinal center (GC) B cells were analyzed by flow cytometry (Table S2) as detailed elsewhere (*43*). Samples were acquired using an LSRFortessa (BD) and analyzed with FlowJo v10.7.1 (Fig. S10) (TreeStar). Antibody secreting cells (ASC) were measured by ELISpot in triplicate as previously reported (*39*) using S-2P protein (1 µg/well) and goat-anti-human IgG (Southern Biotech). Plates were developed using the BD AEC kit. Results are reported as ASC per million input cells.

#### T Cell Responses

PBMC or LN-MNC were stimulated with 2 µg/mL overlapping peptides encompassing the SARS-CoV-2 Spike protein (JPT Technologies) or vehicle DMSO in the presence of co-stimulatory antibodies against CD28 and CD49d and quantified by flow cytometry (Table S3) as described (*43*). Activation-induced marker (AIM) T follicular helper cell (Tfh) responses were quantified in LN-MNC stimulated with 5 µg/mL Staphylococcal enterotoxin B (Toxin Technologies) or vehicle DMSO for 18h (*23*) and analyzed by flow cytometry (Table S3, Fig. S15).

#### Statistical analyses

Spearman’s rank correlations were estimated between pre-specified parameters at specific timepoints. Statistical analyses were performed using SAS version 9.4 (Cary, NC, USA). ID50 neutralization titer decline half-life was estimated based on random-effects regression models of decay with first-order kinetics (*44, 45*). Models were fit separately by vaccine group and data prior to the second vaccine dose (i.e., from weeks 0 and 4) were excluded. Bi-phasic decline was modeled using a linear spline with one knot (*46*), with different knots considered ranging from 8 to 18 weeks. For each vaccine group, the model with the knot at 18 weeks fit the best according to the Akaike Information Criterion.

### Supplementary methods

#### Preparation of the SARS CoV2 Spike protein

The CoV2 Spike (*47*) (2019-nCoV) protein was transiently expressed in Freestyle HEK293 cells (Thermo Scientific) by first diluting the plasmid in Opti-MEM-1 (Gibco) medium; 0.8mg of plasmid in 25mL of Opti-MEM-1 for each liter of cells transfected and sterile filtered using a 0.22 µm filter. 1 mL of 293fectin (Gibco) was diluted in 25 mL of Opti-MEM-1 and allowed to incubate at room temperature for 5±1 minutes. The plasmid and the 293fectin mixtures were combined and swirled to mix, then incubated at room temperature for 25±5 minutes. During the incubation, the Freestyle HEK293 cells were diluted to 1.25×106 cells/ mL in a 2-liter Corning flask. At the end of the 25±5 minute incubation of the 293fectin and plasmid mixtures, 50mL of the transfection mixture was added to each flask of diluted cells with gentle agitation of the flask during addition. The cell + transfection mixture was then split between two 2-liter Corning flasks for a final volume of approximately 500mL each and placed on a platform shaker in a humidified incubator at 37°C with 8% CO2. The shaker speed was set to 120rpm and the flasks allowed to incubate for 6 days. At the end of the 6-day incubation, the flasks containing the transfected cells were removed from the incubator and the contents of the flasks were aseptically transferred to 500mL Corning centrifuge tubes. The supernatant was clarified by centrifugation and subsequently filtered using a 0.8µm filter bottle system. Keeping the clarified and filtered supernatant on ice, the supernatant was concentrated using a Sartorius Vivaflow 200 30kDa TFF system. In preparation for the purification of the CoV2 Spike protein, 4mL of Strep-Tactin Resin (iba Life Science) was placed in a conical tube. The concentrated supernatant was transferred to the conical tube containing the Strep-Tactin resin and allowed to bind with gentle agitation. The supernatant and resin were then transferred to a polypropylene column for gravity-flow chromatography (BioRad). Once the resin had settled in the column the resin was washed and the protein eluted following instructions supplied by the vendor (iba Life Science). The eluate was buffer-exchanged into 2mM Tris and 200mM Sodium Chloride storage buffer using Millipore Centrifugal filters. The resulting material was finally filtered through a 0.2um syringe filter (Pall) and stored in storage buffer. Additional purification by size exclusion chromatography over a Superose 6 column was performed to increase purity.

#### Measurement of S-specific IgG and IgA in saliva

Saliva was collected with absorbent Merocel sponges (Beaver Visitec) by placing a sponge between the cheek and gum in the back of the mouth for 5 minutes. Secretions were eluted by centrifugation at 18,000 g and 4°C after addition of 50µl of PBS containing protease inhibitors (*48*), 1% TritonX-100, 1% BSA, 0.05% azide, and 0.05% Tween-20 to sponges. A customized binding antigen multiplex assay (BAMA) was used to measure IgG or IgA antibodies to SARS CoV-2 recombinant receptor binding domain protein (RBD; generously provided by Dr. Wrammert, Emory University, Atlanta, GA) and S2 extracellular domain (SinoBiologicals #40590-V08B, Wayne, PA). Briefly, proteins were dialyzed in PBS and conjugated to Bioplex Pro carboxylated magnetic beads (Bio-Rad, Hercules, CA) using N-hydroxysulfosuccinimide and ethylcarbodiimide as described (*49*). Serial dilutions of standard and centrifuged salivary secretions in PBS containing 1% TritonX-100, 1% BSA, 0.05% azide, and 0.05% Tween-20 were mixed with RBD+S2 beads overnight at 1100rpm and 4°C using a plate mixer. The IgG standard was a cocktail of anti-S1 RBD (Genscript #HC2001) and anti-S2 (SinoBiologicals #40590-D001) humanized IgG monoclonal antibodies. The standard for IgA assays was a pooled serum from infected rhesus macaques (*50*) that had been calibrated relative to the previously mentioned monoclonal antibodies. The following day, beads were alternately washed using a Bio-Rad BioPlex wash station and treated for 30min with 2µg/ml biotinylated affinity-purified goat antibody to human γ chain (SouthernBiotech Associates, Birmingham, AL) or clone IgA5-3B mouse anti-monkey IgA (Bio-Rad) followed by 1/400 avidin-phycoerythrin (Southern Biotechnology Associates). A Bio-Rad Bioplex 200 and BioManager software were used to measure fluorescent intensity and construct standard curves for interpolation of antibody concentrations in test samples. Concentrations of antibody measured in BAMA were normalized relative to the total IgG and IgA measured by ELISA as described (*48*) using plates coated with goat anti-monkey IgG or IgA and the secondary antibodies above.

#### Mapping plasma S-specific IgG

SARS-CoV-2 antigens, including whole spike (produced by PPF), S1 (Sinobiological cat# 40591-V08H), S2 (Sinobiological cat# 40590-V08B), RBD (Sinobiological cat# 40592-V08H) and NTD (Sinobiological cat# 40591-V49H) were conjugated to Magplex beads (Bio-Rad, Hercules, CA). The conjugated beads were incubated on filter plates (Millipore, Stafford, VA) for 30 min before plasma samples were added. Plasma samples were diluted in assay diluent (1% dry milk, 5% goat serum, and 0.05% Tween 20 in 1× phosphate buffered saline, pH 7.4.) at a 1:10,000-point dilution. Beads and diluted samples were incubated for 30 min with gentle rotation, and IgG binding was detected using a phycoerythrin (PE)-conjugated mouse anti-monkey IgG (Southern Biotech, Birmingham, Alabama) at 2 μg/ml. Plates were washed and acquired on a Bio-Plex 200 instrument (Bio-Rad, Hercules, CA), and IgG binding was reported as mean fluorescence intensity (MFI). To assess assay background, the MFIs of wells without sample (blank wells) were used, as well as evaluated nonspecific binding of the samples to unconjugated blank beads.

#### SARS-CoV-2 Pseudovirus Neutralization Assay

SARS-CoV-2 neutralization was assessed with Spike-pseudotyped viruses in 293T/ACE2 cells as a function of reductions in luciferase (Luc) reporter activity. 293T/ACE2 cells were kindly provided by Drs. Farzan and Mu at Scripps Florida. Cells were maintained in DMEM containing 10% FBS, 25 mM HEPES, 50 µg/ml gentamycin and 3 µg/ml puromycin. An expression plasmid encoding codon-optimized full-length Spike of the Wuhan-1 strain (VRC7480), was provided by Drs. Graham and Corbett at the Vaccine Research Center, National Institutes of Health (USA). The D614G amino acid change was introduced into VRC7480 by site-directed mutagenesis using the QuikChange Lightning Site-Directed Mutagenesis Kit from Agilent Technologies (Catalog # 210518). The mutation was confirmed by full-length Spike gene sequencing. Pseudovirions were produced in HEK 293T/17 cells (ATCC cat. no. CRL-11268) by transfection using Fugene 6 (Promega Cat#E2692) and a combination of Spike plasmid, lentiviral backbone plasmid (pCMV ΔR8.2) and firefly Luc reporter gene plasmid (pHR’ CMV Luc) (*51*) in a 1:17:17 ratio. Transfections were allowed to proceed for 16-20 hours at 37oC. Medium was removed, monolayers rinsed with growth medium, and 15 ml of fresh growth medium added. Pseudovirus-containing culture medium was collected after an additional 2 days of incubation and was clarified of cells by low-speed centrifugation and 0.45 µm micron filtration and stored in aliquots at −80°C. TCID50 assays were performed on thawed aliquots to determine the infectious dose for neutralization assays.

For neutralization, a pre-titrated dose of virus was incubated with 8 serial 5-fold dilutions of serum samples in duplicate in a total volume of 150 µl for 1 hr at 37oC in 96-well flat-bottom poly-L-lysine-coated culture plates (Corning Biocoat). Cells were suspended using TrypLE Select Enzyme solution (Thermo Fisher Scientific) and immediately added to all wells (10,000 cells in 100 µL of growth medium per well). One set of 8 control wells received cells + virus (virus control) and another set of 8 wells received cells only (background control). After 66-72 hrs of incubation, medium was removed by gentle aspiration and 30 µL of Promega 1X lysis buffer was added to all wells. After a 10-minute incubation at room temperature, 100 µl of Bright-Glo luciferase reagent was added to all wells. After 1-2 minutes, 110 µl of the cell lysate was transferred to a black/white plate (Perkin-Elmer). Luminescence was measured using a PerkinElmer Life Sciences, Model Victor2 luminometer. Neutralization titers are the serum dilution at which relative luminescence units (RLU) were reduced by either 50% (ID50) or 80% (ID80) compared to virus control wells after subtraction of background RLUs. Serum samples were heat-inactivated for 30 minutes at 56C prior to assay.

#### Whole virus neutralization assay

Neutralization of SARS-CoV-2 nanoLUC carrying the D614G mutation was assessed as described in Hou et al with modifications (*52*). Briefly, under BSL-3 containment, serially diluted sera at 8 dilutions were incubated for one hour with SARS-CoV-2 D614G nanoLUC virus at 5% CO2 and 37°C. After incubation, the virus/antibody mixtures were added in duplicate to black 96-well plates containing Vero E6 cells (2 x 104 cells/well). Each plate contains virus-only (no serum) control wells. The plates were incubated for 24 hr at 37 °C, 5% CO2, the cells lysed, and luciferase activity measured with the Nano-Glo Luciferase Assay System (Promega). Neutralization activity is expressed as the dilution concentration at which the observed relative light units (RLU) are reduced by 50% or 80% relative to virus-only control wells.

#### Plasma ACE2 blocking assay

Corning 384-well plates were coated with 3.5 ug/mL ACE2 protein (Sinobiological) 24 hours before conducting the experiment. The day of the assay, the plate was blocked with assay diluent (phosphate-buffered saline containing 4% whey, 15% normal goat serum, and 0.5% Tween 20) for 1 hour at room temperature. Plasma was diluted 1:10, 1:40, or 1:60 using the assay diluent, and incubated with 1 ug/mL HRP-RBD protein (Genescript) at 37C for 1 hour. Plates were washed and the preincubated mixture of plasma and HRP-RBD protein was added in duplicates and incubated at RT for 1 hour. Then, plates were washed 4 times to remove unbound sample, and peroxidase substrate solution (SeraCare) was added for 4 minutes before stopping the reaction using stop solution (SeraCare). Signal was detected using a Spectramax plate reader at OD 450. The amount of signal detected in wells without sample (diluent only) was considered the maximal binding response, and the OD450 detected in the sample wells was transformed to percentage inhibition compared to maximum binding. Data presented is the average of two replicates when CV is less than 20%, and if different assays were performed, median value was considered.

#### Preparation of antigen-specific hook reagents for flow cytometry

To express the prefusion S ectodomain, a gene encoding residues 1−1208 of 2019-nCoV S (GenBank: MN908947) with proline substitutions at residues 986 and 987, a “GSAS” substitution at the furin cleavage site (residues 682–685), a C-terminal T4 fibritin trimerization motif, an HRV3C protease cleavage site, a TwinStrepTag and an 8XHisTag was synthesized and cloned into the mammalian expression vector pαH. To express the 2019-nCoV RBD-SD1, residues 319−591 of 2019-nCoV S were cloned upstream of a C-terminal HRV3C protease cleavage site, a monomeric Fc tag and an 8XHisTag. Similarly, to express the SARS-CoV RBD-SD1, residues 306−577 of SARS-CoV S (Tor2 strain) were cloned upstream of a C-terminal HRV3C protease cleavage site, a monomeric Fc tag and an 8XHisTag. Dimers are prepared based on the molar ratio (2:1) of the analyte protein and fluorochrome-conjugated Strep-Tactin, respectively. PE (IBA GmbH, 6-5000-001) or APC (IBA GmbH, 6-5010-001) conjugated Strep-Tactin was reacted with the protein over 5 additions, incubating for 15 minutes between each addition. The final concentration of tetramer was calculated with respect to the analyte protein. The solution was aliquoted based on the expected usage per experiment, snap frozen, and stored at −80°C. For quality control, monoclonal antibodies were bound to polystyrene beads (Spherotech) per the manufacturer’s instructions. B cell hooks were tested on beads coated with Ab026204 and CH65; mAb Ab026204 binds SARS-CoV2, mAb CH65 binds influenza hemagglutinin (*53*) and was used a negative control. Briefly, 0.5 µL of beads were diluted to 25 µL in PBS+0.02% NaN3. The bead mixture was added to 6 wells (25 µL per well) in a 96-well filter plate. B-cell hooks were diluted to a final concentration of 40 µg/mL in 175 µL of PBS+0.02% NaN3, then four serial 2-fold dilutions were made down to a final concentration of 5 µg/mL. Test conjugate dilutions (25 µL per well) were added to each bead set, with one additional well of each bead set left as an unstained control. After 30-minute incubation, the solution was vacuumed off and the beads washed 3 times with 150 µL PBS+0.02% NaN3. The washed beads were resuspended in 125 µL of PBS+0.02% NaN3, and the samples were run on a BD LSRII cytometer within 4 hours of preparation. Data were analyzed using Flow Jo software (Treestar) and plots of each dilution compared to the unstained control.

#### Antigen-specific B cell quantitation

Freshly isolated or archived (week 6) PBMC or LNC (2×106 cells) were washed with phosphate buffered saline (PBS, Gibco) and pelleted by centrifugation at 500 x g for 7 minutes. Supernatant was discarded and cells were resuspended in 150 µL 1% bovine serum albumin (BSA, Sigma-Aldrich) + 5 µM Chk2 inhibitor II in PBS and incubated 15 minutes at 4C in the dark. Cells were subsequently washed with PBS + 1% BSA (7 minutes at 500 x g). The cell pellet was stained with antibodies prescribed in Table S2 for 30 minutes in the dark at 4C. After staining, cells were washed as before and fixed with 1% paraformaldehyde before immediate acquisition on an LSRFortessa (BD) using BD FACSDiva v.8.0 and analyzed with FLowJo software v10.7.1 (TreeStar).

#### B Cell ELISpot

Polyvinylidene fluoride membranes (Millipore) were activated with 70% ethanol for 1 minute, subsequently coated with 1 μg SARS-CoV-2 Spike protein (2019-nCoV) per well and blocked for 2 hours with 2% milk and stored at 4C. Fresh or thawed cell preparations were washed and stimulated (106 cells/mL) with 1 μg/mL R848 (InvivoGen, San Diego, CA) and 10 ng/mL human IL-2 (Miltenyi Biotec) for 72 hours in complete medium (cRPMI): RPMI-1640 medium (Gibco) containing penicillin, streptomycin, L-glutamine (Sigma-Aldrich), and 10% heat-inactivated FBS (Gibco). Stimulated cells (8 × 104/well) were incubated in triplicate wells of SARS-CoV-2 Spike protein-coated microtiter plates overnight at 37°C, washed with PBS+0.05% Tween 20, and incubated 1 hour with 1 μg/mL biotinylated affinity-purified goat anti-human IgG (Southern Biotech) at room temperature. Plates were washed with PBS+0.05% Tween 20, incubated with a 1:4000 dilution of avidin-peroxidase (Southern Biotech) for 1 hour at room temperature, and developed using the BD AEC kit using 100 µL per well. Dried membranes were analyzed with an automated ELISpot Reader System (Autoimmun Diagnostika GmbH). Results are reported as the number of antibody-secreting cells (ASC) per 106 MNCs.

#### T cell cytokine assay

Frozen cells were thawed, or fresh preparations were washed and cultured in cRPMI (106/mL) with or without 2 µg/mL overlapping peptides spanning the length of SARS-CoV-2 spike protein (JPT Technologies) or DMSO vehicle together with antibodies against co-stimulatory CD28 and CD49d from BD. Naïve donor cells were included with each assay and, in addition to peptide pool or vehicle, were stimulated with 0.5x cell stimulation cocktail (eBiosciences) as a positive control. Cells were surface stained as outlined in Table S2 and then permeabilized with BD CytoFix/CytoPerm per manufacturer’s recommendations and stained with intracellular antibodies as prescribed in Table S2. Data were collected using an LSRFortessa and BD FACSDiva v8.0 and analyzed with FlowJo software v10.7.1 (TreeStar). Intracellular cytokine gates were Boolean gated and reported as single positive events unless otherwise noted.

#### Activation-induced marker (AIM) Assay

The activation-induced marker (AIM) assay was performed on cryopreserved LNC. Cells were thawed, rested 3 h at 37°C with 5% CO2, resuspended in AIM V medium (Gibco), and transferred at 106 cells per well to a 24-well plate. Cells were cultured with vehicle DMSO (negative control) or with 0.5 μg/ml staphylococcal enterotoxin B (Toxin Technologies) for 20 hours at 37°C with 5% CO2. After stimulation, cells were stained with prescribed antibodies per Table S2 and acquired immediately on an LSRFortessa instrument running FACSDiva v8.0 software (BD Biosciences) and analyzed as described above.

### Supplementary figures

**Figure S1:**
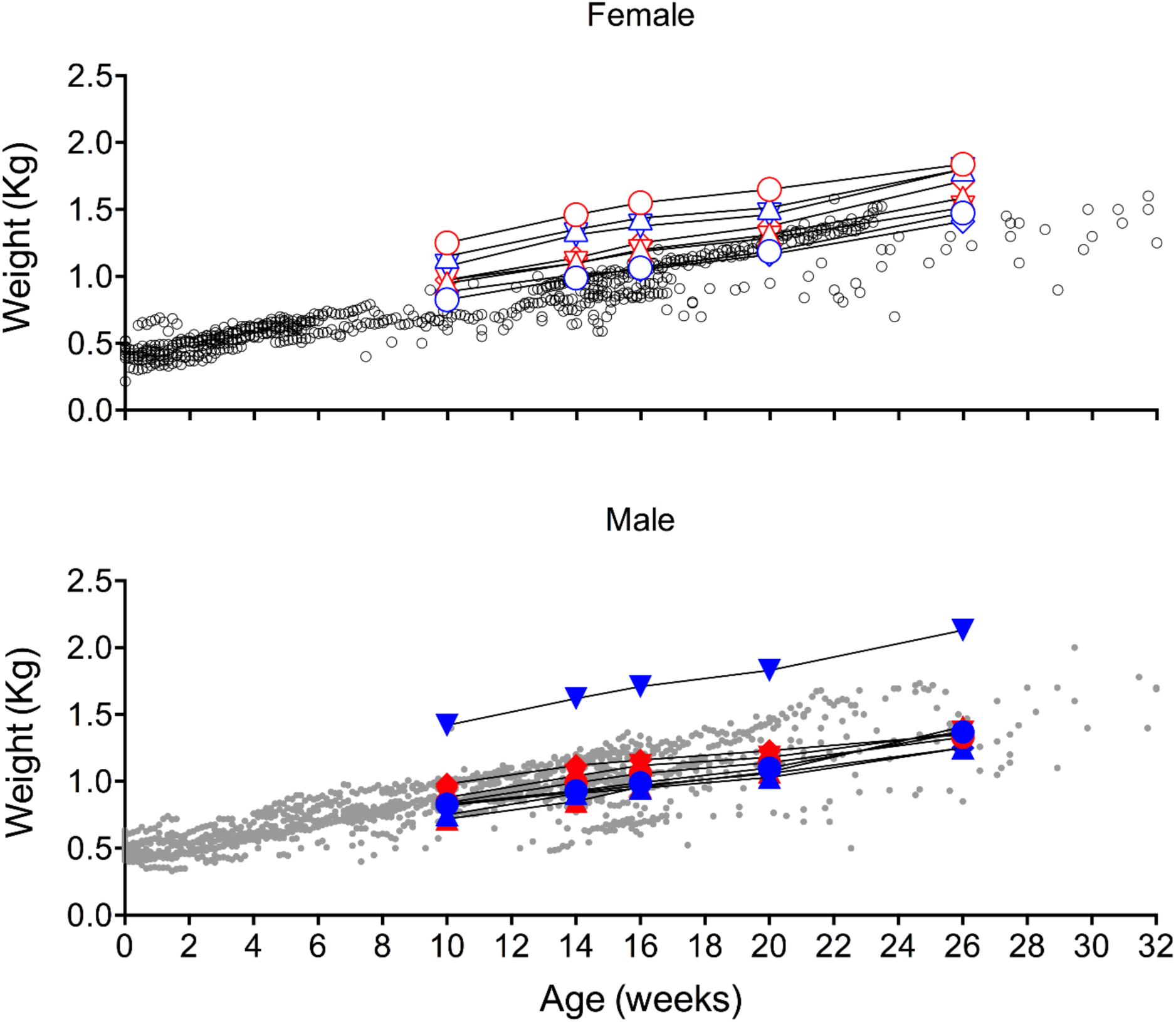
Weight gain of SARS-CoV-2 immunized infant rhesus macaques. Longitudinal weight data for all female (top) and male (bottom) animals from this study with their respective symbol shapes and colors (Table S1) overlaid with historical weight data from age and sex-matched infant rhesus monkeys housed outdoors at the CNPRC, Davis California (open black circles-female; filled gray circles-male).

**Figure S2.**
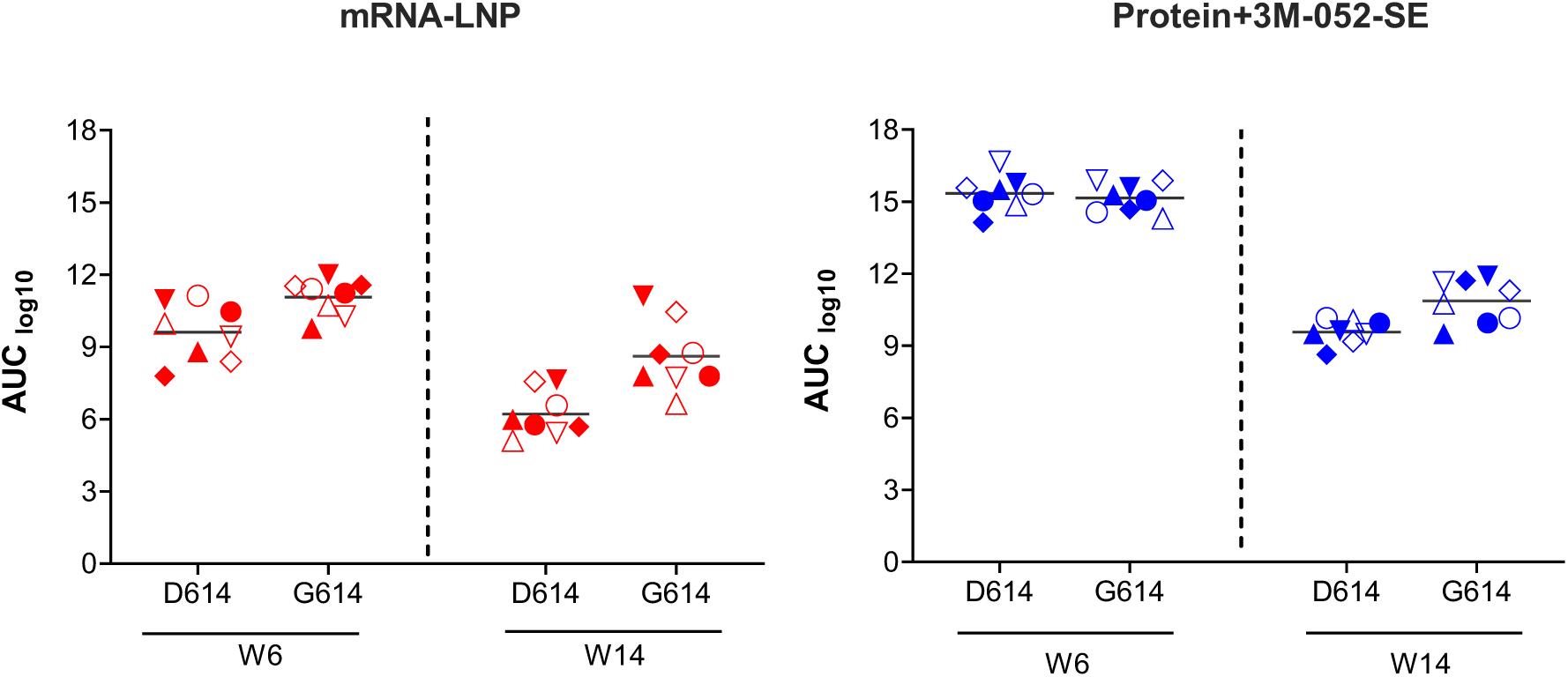
IgG binding to the D614 spike protein and the D614G variant. Plasma IgG binding response was assessed by ELISA using serial dilutions of plasma collected at W6 and W14. Area under the curve (AUC) was compared between binding to D614 spike protein, the immunogen in both vaccines, and the G614 spike protein that was used in the neutralization assays because it represented the dominant variant at study initiation. Different symbols represent individual animals in the mRNA-LNP (red) or Protein+3M-052-SE (blue) vaccine groups, respectively (Table S1). Horizontal lines represent median values.

**Figure S3:**
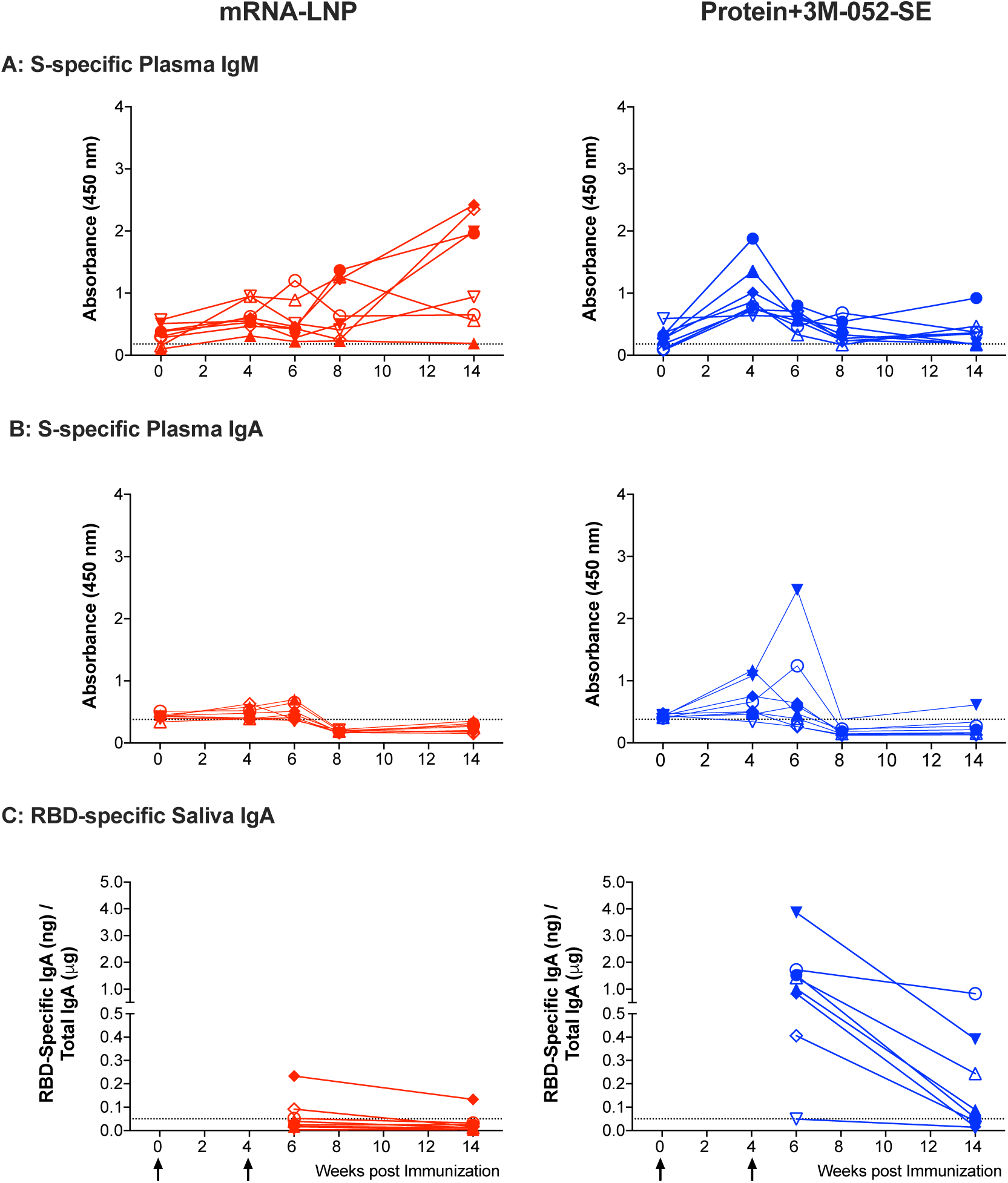
Plasma IgM and IgA and salivary IgA responses in infant rhesus macaques. **Panels A and B** illustrate longitudinal plasma S-specific IgM (Panel A) and IgA (Panel B) levels in 1:10 diluted plasma samples, reported as absorbance readings at 450 nm. In **Panel C**, we report receptor-binding domain-specific salivary IgA concentrations in ng per µg of total IgA. Note that salivary IgA was only measured at weeks 6 and 14. Red or blue lines and symbols represent the mRNA or protein vaccine groups, respectively, with different symbols representing individual animals (Table S1).

**Figure S4.**
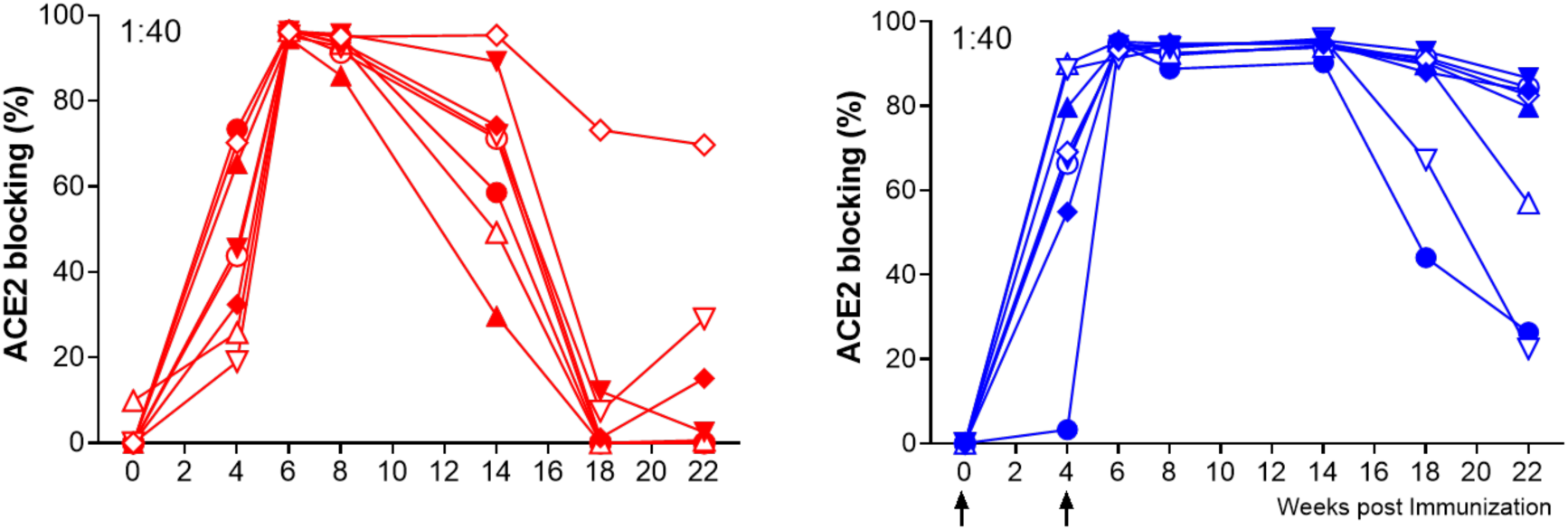
Plasma ACE2 blocking responses. ACE2 blocking assay performed at 1:40 of vaccinated animals in the mRNA-LNP (red) or Protein+3M-052-SE vaccine (blue) groups over time, with different symbols representing individual animals (Table S1).

**Figure S5.**
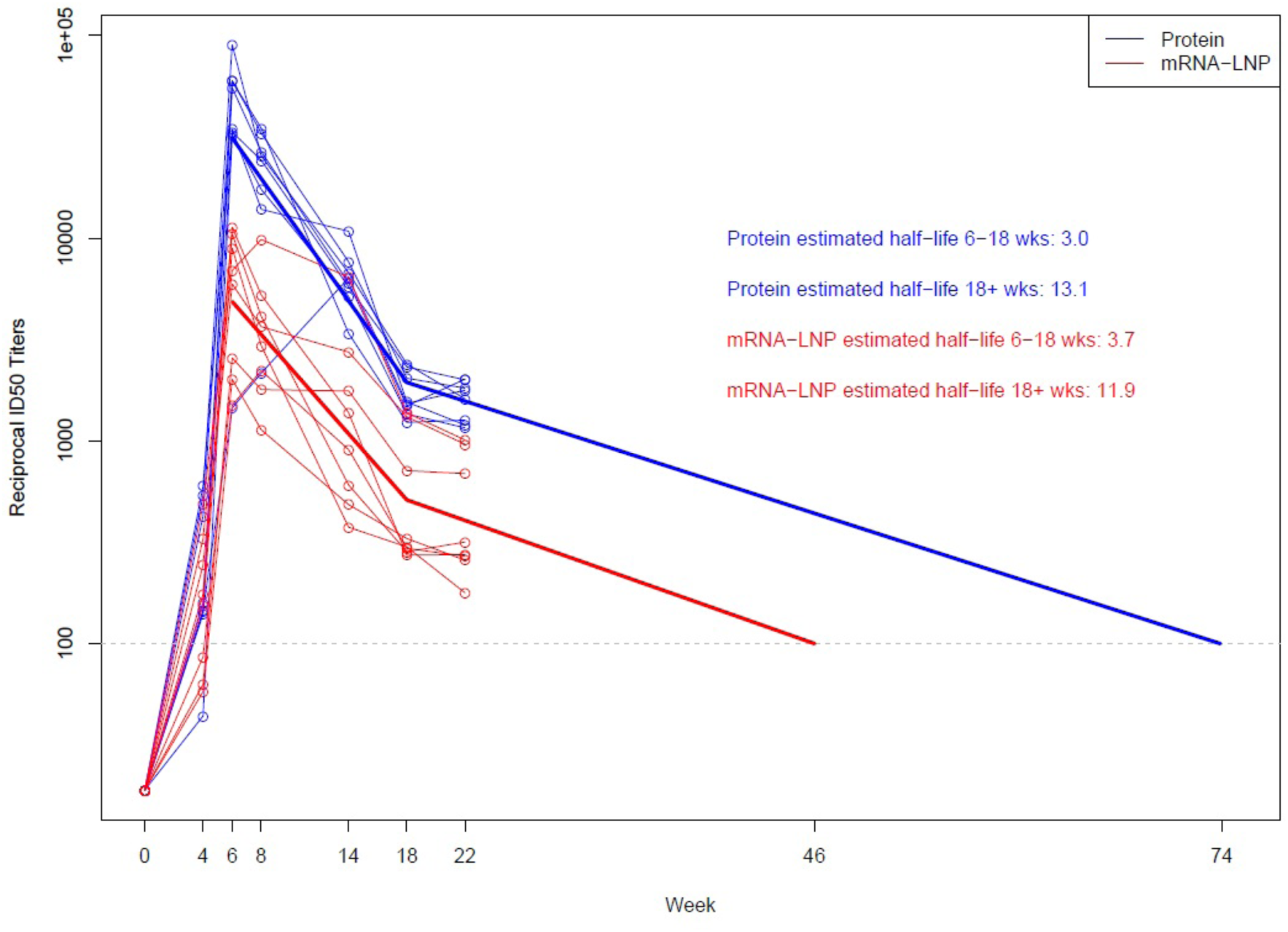
Modelled bi-phasic decline of neutralizing ID_50_ titers. ID50 neutralization titer (obtained using the pseudovirus neutralization assay) decline half-life was estimated based on random-effects regression models of decay with first-order kinetics. Models were fit separately by vaccine group and data prior to the second vaccine dose (i.e., from weeks 0 and 4) were excluded. Bi-phasic decline was modeled using a linear spline with one knot, with different knots considered ranging from 8 to 18 weeks. For each vaccine group, the model with the knot at 18 weeks fit the best according to the Akaike Information Criterion.

**Figure S6.**
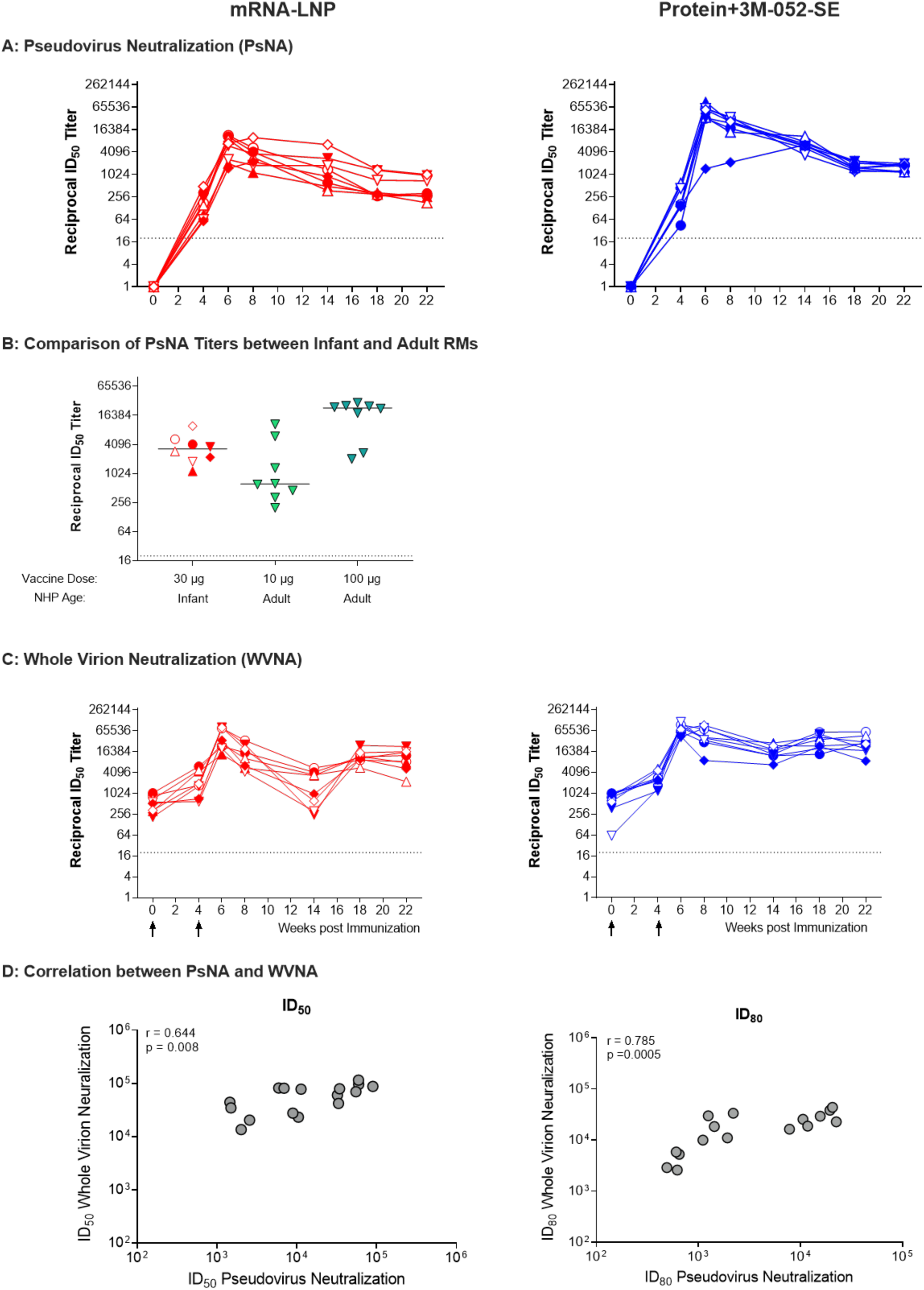
Vaccine-elicited SARS-CoV-2 neutralization responses. **Panel A** shows the ID_50_ neutralization titers obtained using the pseudovirus neutralization assay. **Panel B** depicts the ID_50_ neutralization titers in infant and adult rhesus macaques vaccinated with the mRNA-LNP vaccine (*16*). Infant RM were vaccinated with 30 µg of mRNA-LNP at week 0 and at week 4. Adult RM followed the same vaccine schedule but received 10 µg or 100 µg of the mRNA-LNP vaccine. Neutralization was assessed with a pseudovirus assay at week 8 (4 weeks after the second vaccination). **Panel C** shows the ID_50_ neutralization titers obtained using the whole virus neutralization assay. **Panel D** illustrates the correlation of ID_50_ (left graph) and ID_80_ (right graph) neutralizing titers from both vaccine groups obtained in the pseudovirus or whole virus neutralization assay at week 6. Correlation was assessed using Spearman rank test.

**Figure S7.**
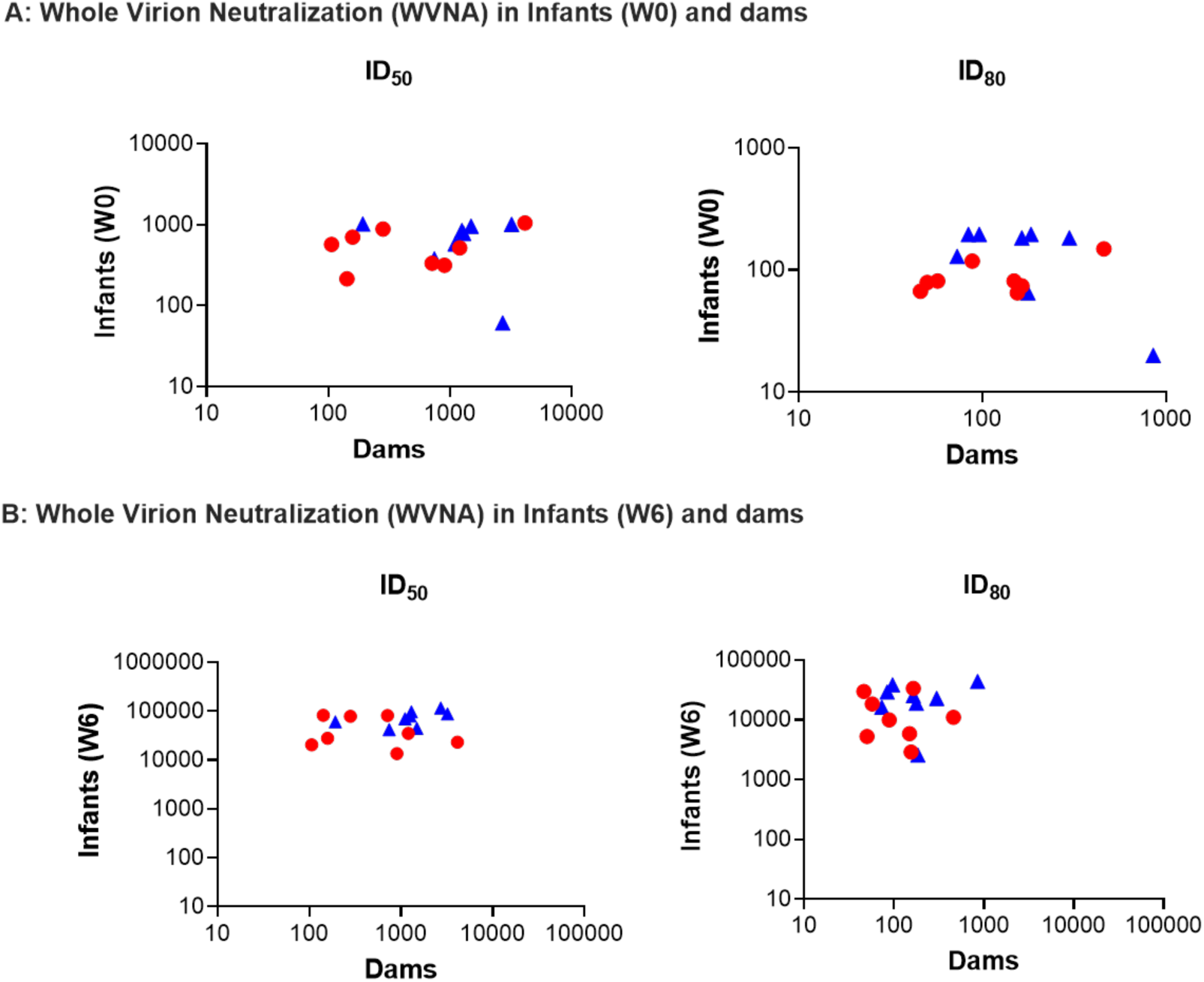
Correlation of whole virion neutralization in dams and infants. Whole virion neutralization (WVNA) was measured in dams and in infants before the vaccine (W0) and after the vaccine boost (W6). There was no correlation of ID_50_ or ID_80_ neutralization titers between infants and dams (Spearman rank analysis). Blue symbols correspond to protein vaccine and red symbols to mRNA-LNP.

**Figure S8.**
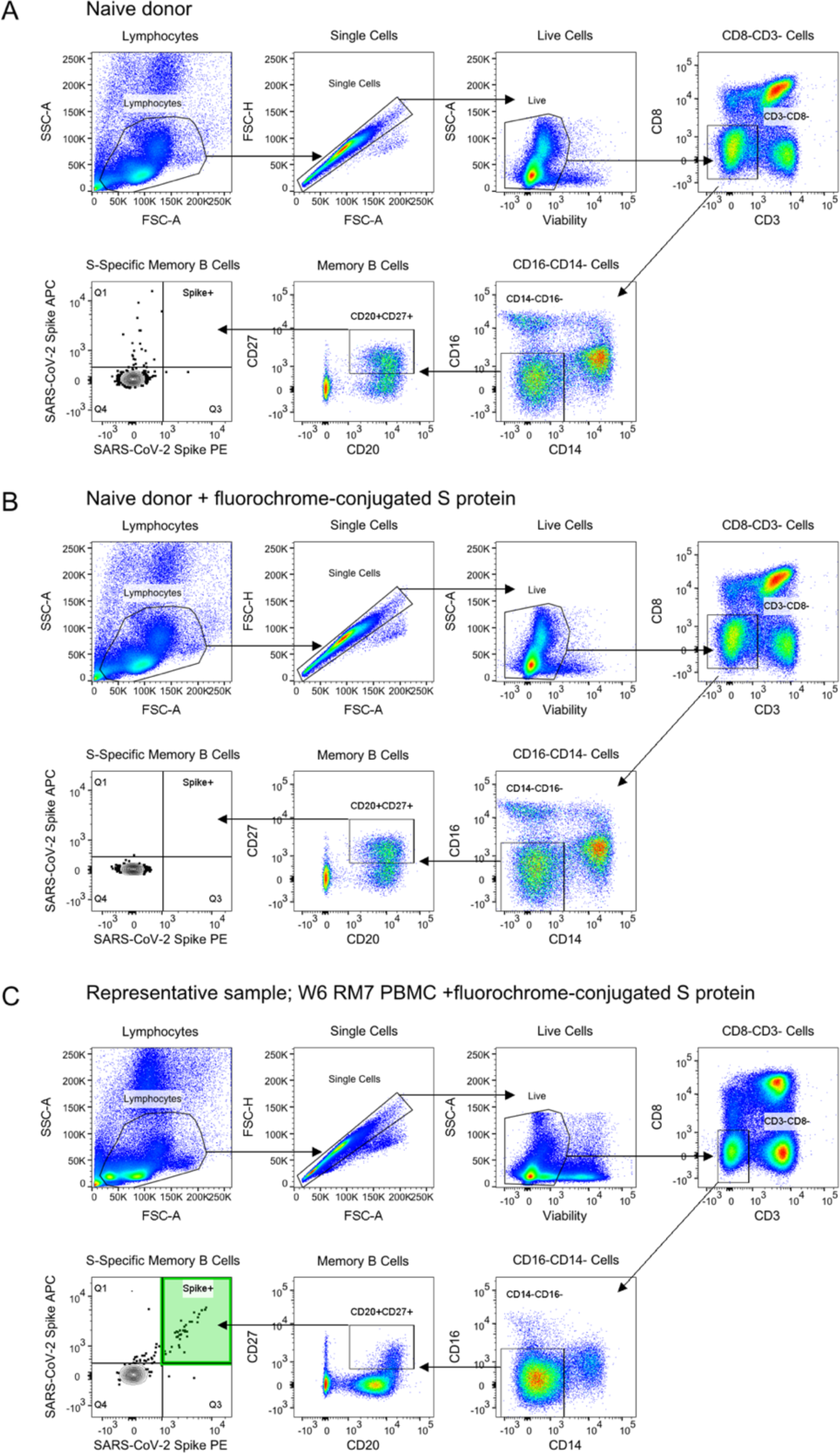
S-specific B cell gating strategy. **Panel A** shows a representative plot from RM7; **Panel B** shows a naive, age-matched control macaque without SARS-CoV-2 S protein conjugates; **Panel C** shows the same donor macaque with the SARS-CoV-2 S protein conjugates included.

**Figure S9.**
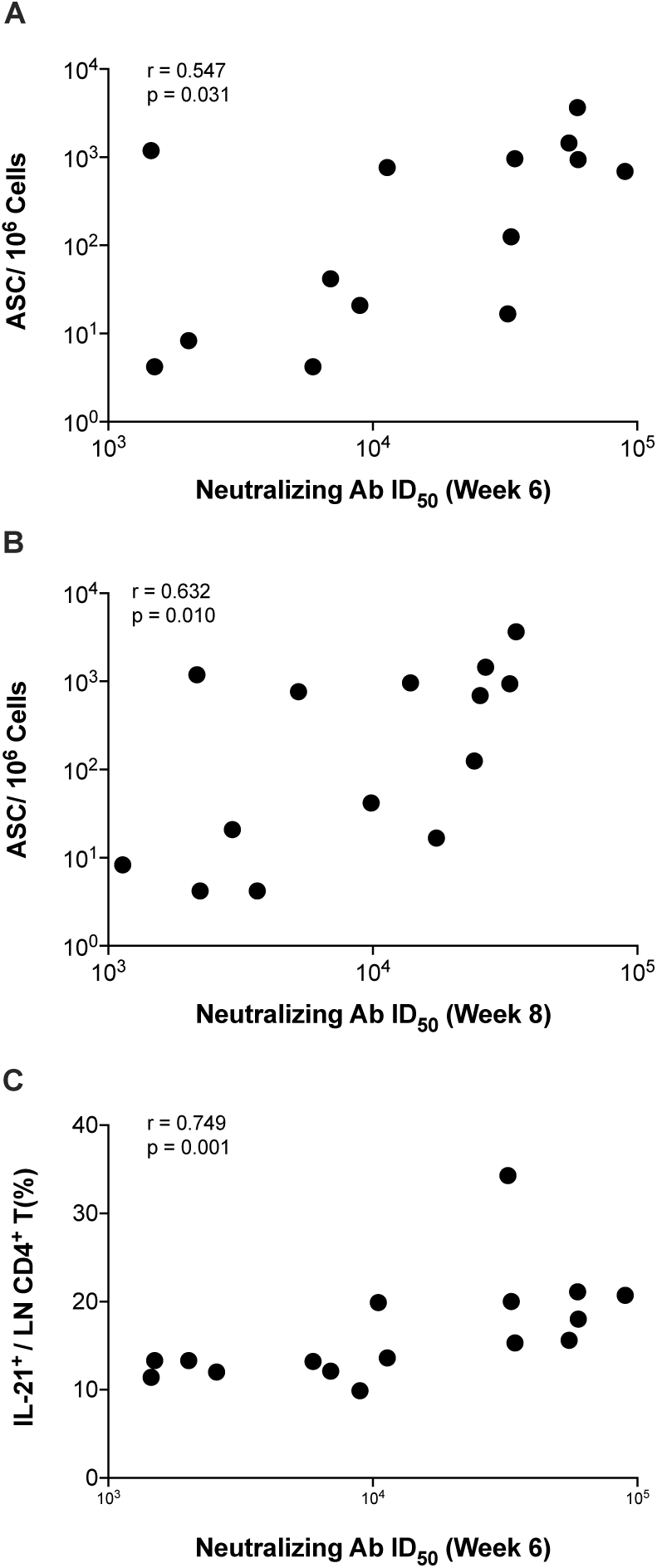
Correlation between neutralizing antibody titers and cellular responses. **Panels A and B** show the correlation between neutralizing antibody ID_50_ titers with lymph node ASC numbers at W6 and W8, respectively. **Panel C** defines the association between lymph node CD4^+^IL-21^+^ T cells at W6 and neutralization titers. Each symbol represents a single animal, animals from both vaccine groups were included in the analysis. Correlations were assessed by Spearman rank test.

**Figure S10.**
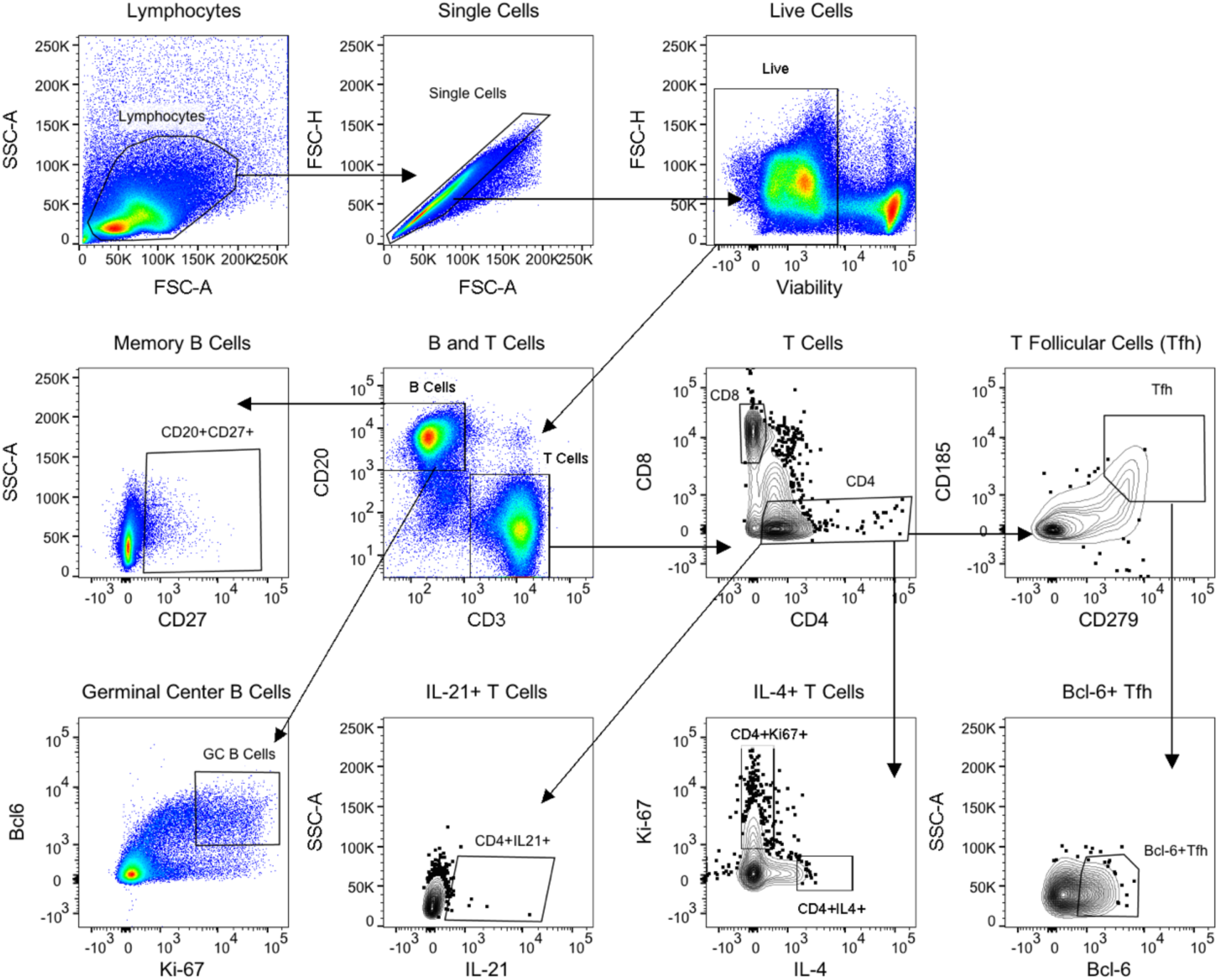
Gating strategy for lymph node germinal center B cells and T cell populations.

**Figure S11:**
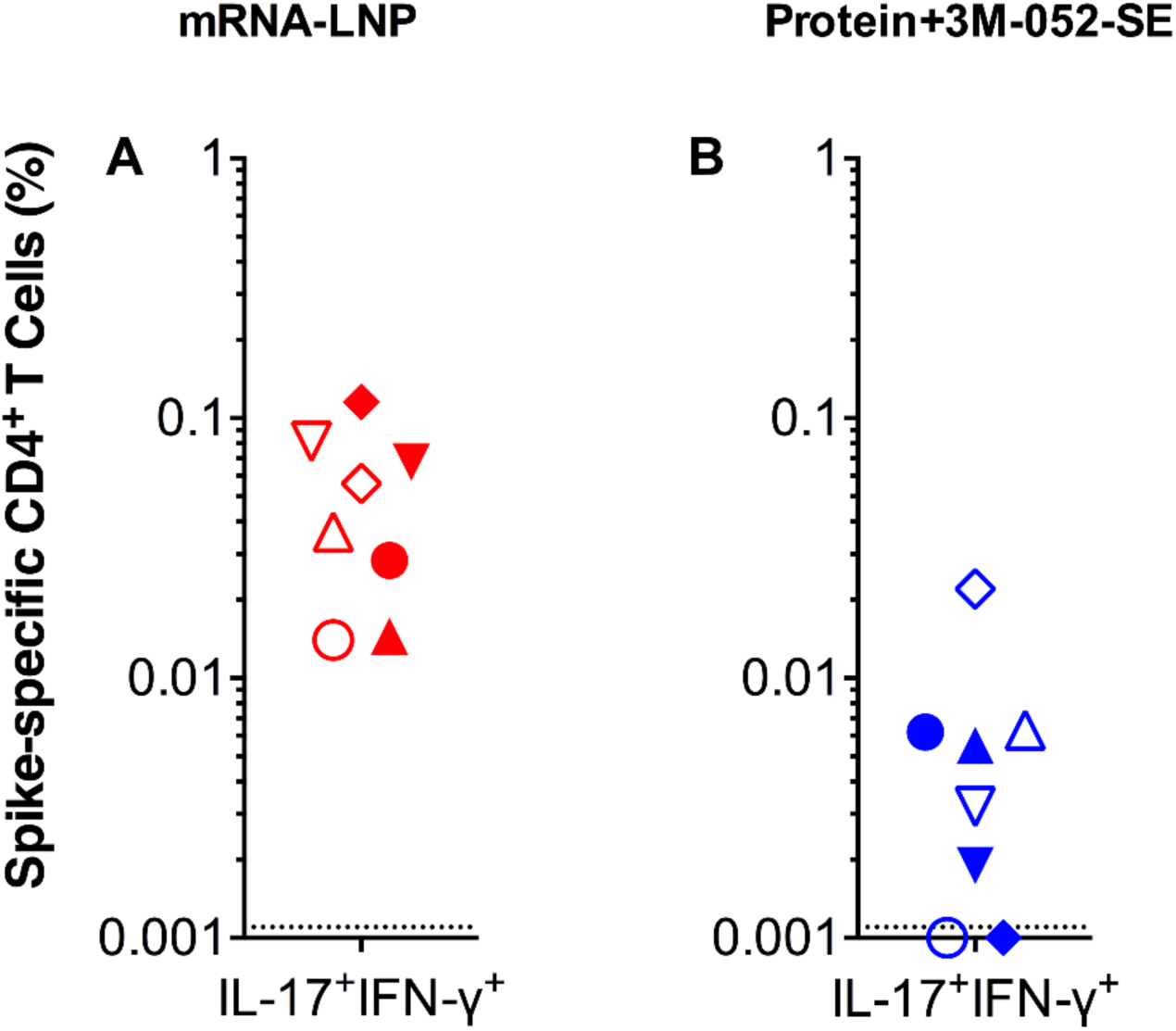
Polyfunctional CD4^+^ T cells in SARS-CoV-2 immunized rhesus macaques at week 14. **Panels A and B** show the frequencies of CD4^+^ T cells that co-produced IL-17 and IFN-γ in response to stimulation with spike protein overlapping peptides in animals of the mRNA-LNP or Protein-3M-052-SE vaccine group, respectively. Individual symbols represent individual animals in each group (Table S1).

**Figure S12.**
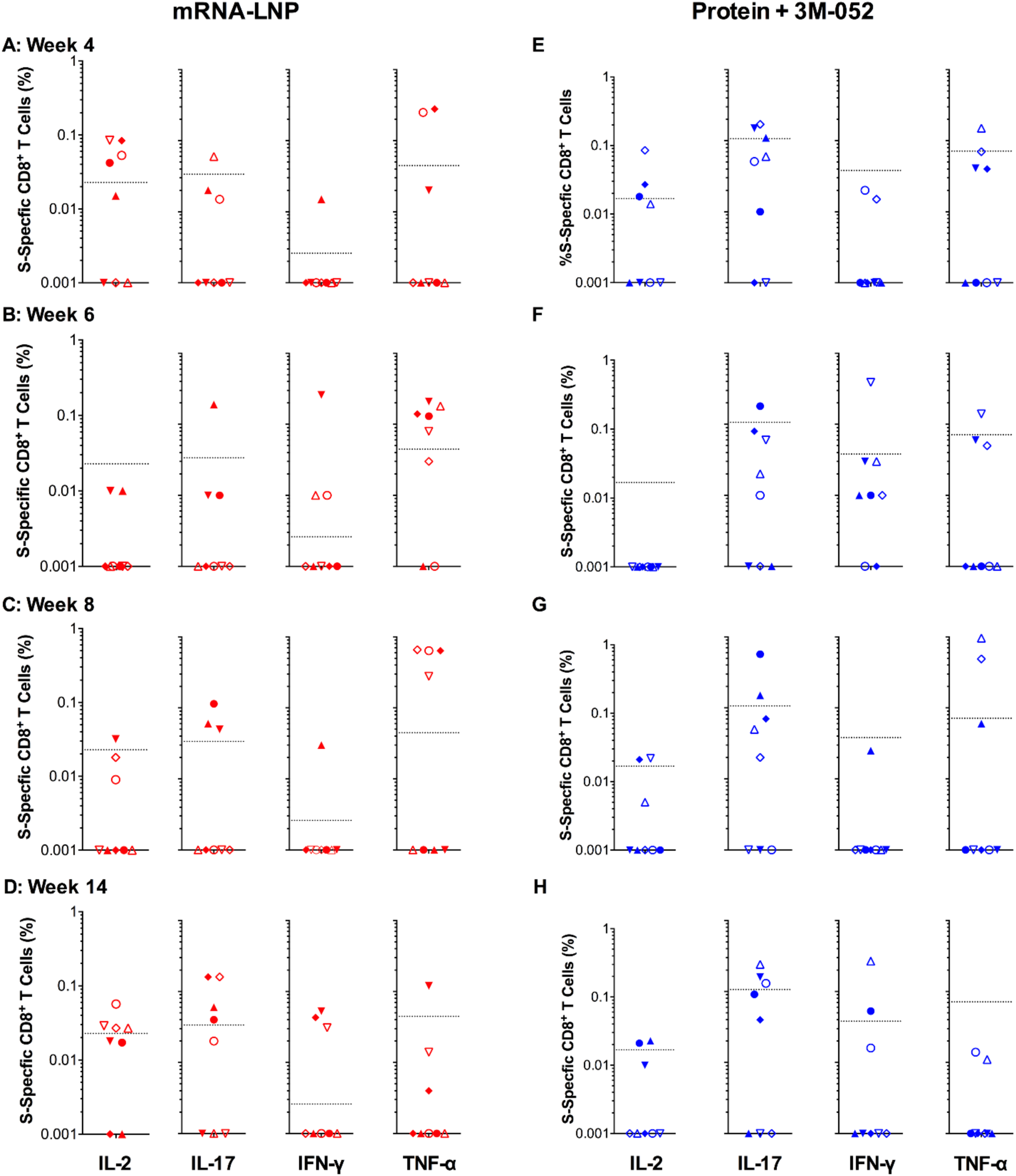
Spike-specific CD8^+^ T cell responses in SARS-CoV-2 immunized infant macaques. Intracellular cytokine staining was performed as described in Figure 5 at weeks 0, 6, 4, 8, and 14 to assess CD8^+^ T-cell responses. **Panels A, B, C and D** show responses detected in PBMC from the mRNA-LNP group at weeks 4, 6, 8, and 14, respectively. **Panels E, F, G and H** show responses in Protein+3M-052+SE vaccinees at weeks 4, 6, 8 and 14, respectively. The legend and symbols used mirror those from Figure 6 (see also Table S1). Dotted lines represent the cut-off for cytokine-positive responses, determined as 2 standard deviations above median values of SARS-CoV-2 naïve animals.

**Figure S13.**
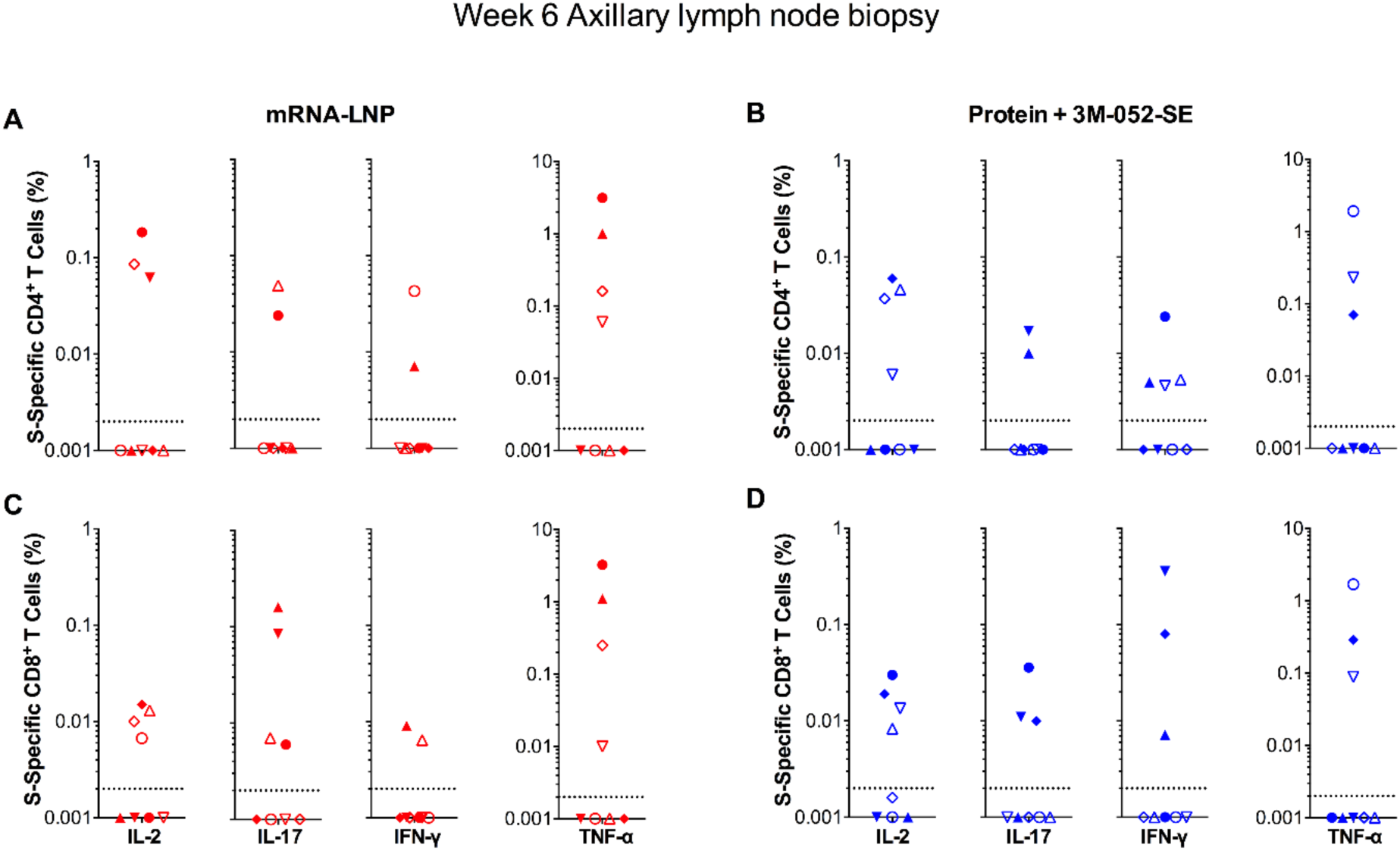
Spike-specific T cell responses in LN following SARS-CoV-2 vaccination. Intracellular cytokine staining as in Figure 5 in lymph node biopsy samples at week 6 to measure Spike-specific T cell responses. **Panels A and B** show mRNA-LNP and Protein+3M-052+SE CD4^+^ T-cell responses, respectively. **Panels C and D** represent CD8^+^ T cell cytokine responses from mRNA-LNP and Protein+3M-052+SE animals, respectively. Symbols and the legend mirror those from Figure 6 (see also Table S1). Dotted lines represent 2 standard deviations from SARS-CoV-2 naïve LN samples.

**Figure S14.**
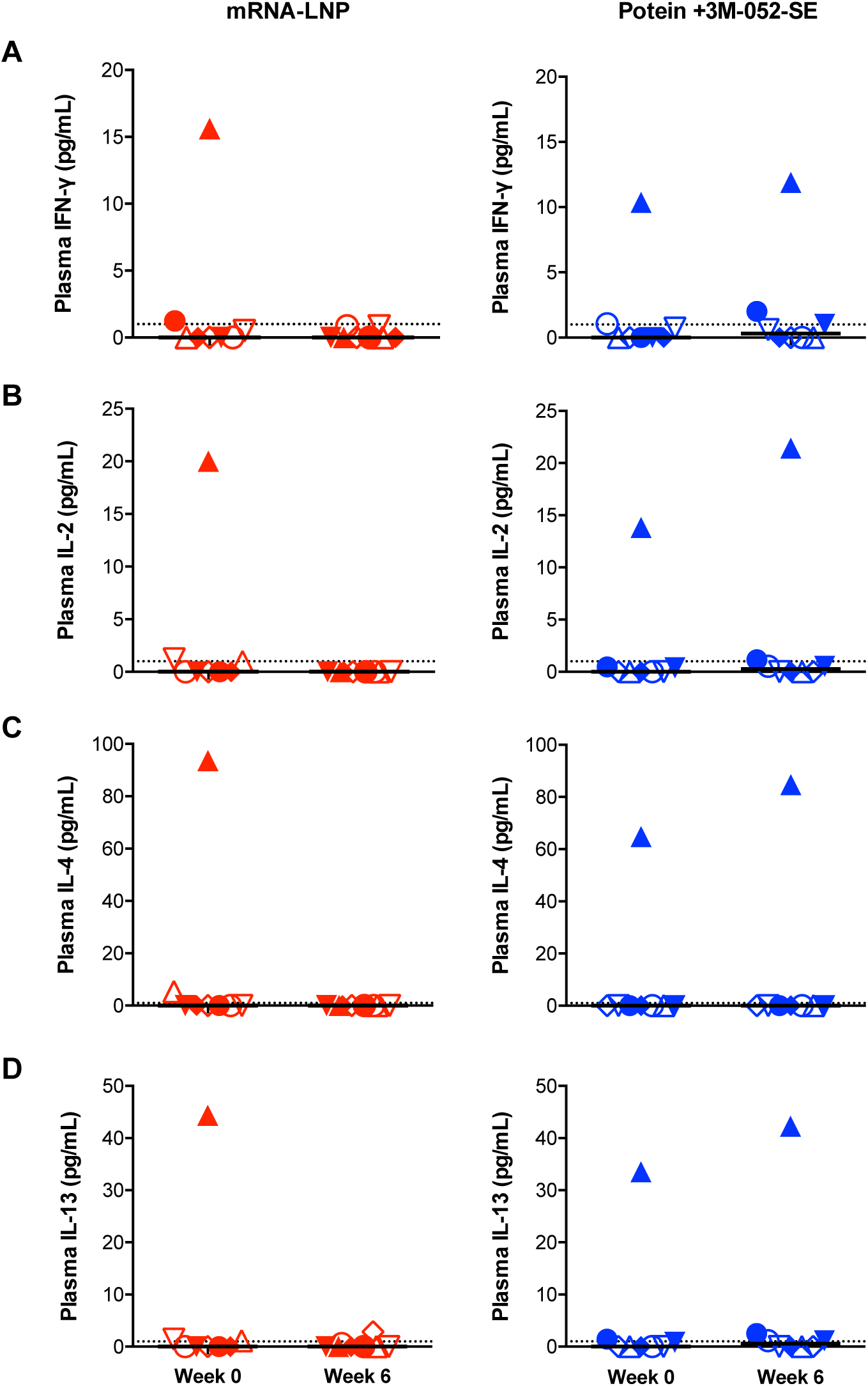
Th1 and Th2 cytokines in plasma of infant rhesus macaques prior to and following SARS-CoV-2 vaccination. Plasma levels of IFN-γ (**panel A**) IL-2 (**panel B**), IL-4 (**panel C**) and IL-13 (**panel D**) were measured by multiplex Luminex assay. Different symbols represent individual animals in the mRNA-LNP (red) or Protein+3M-052-SE (blue) vaccine groups, respectively (Table S1). Horizontal lines represent median values.

**Figure S15.**
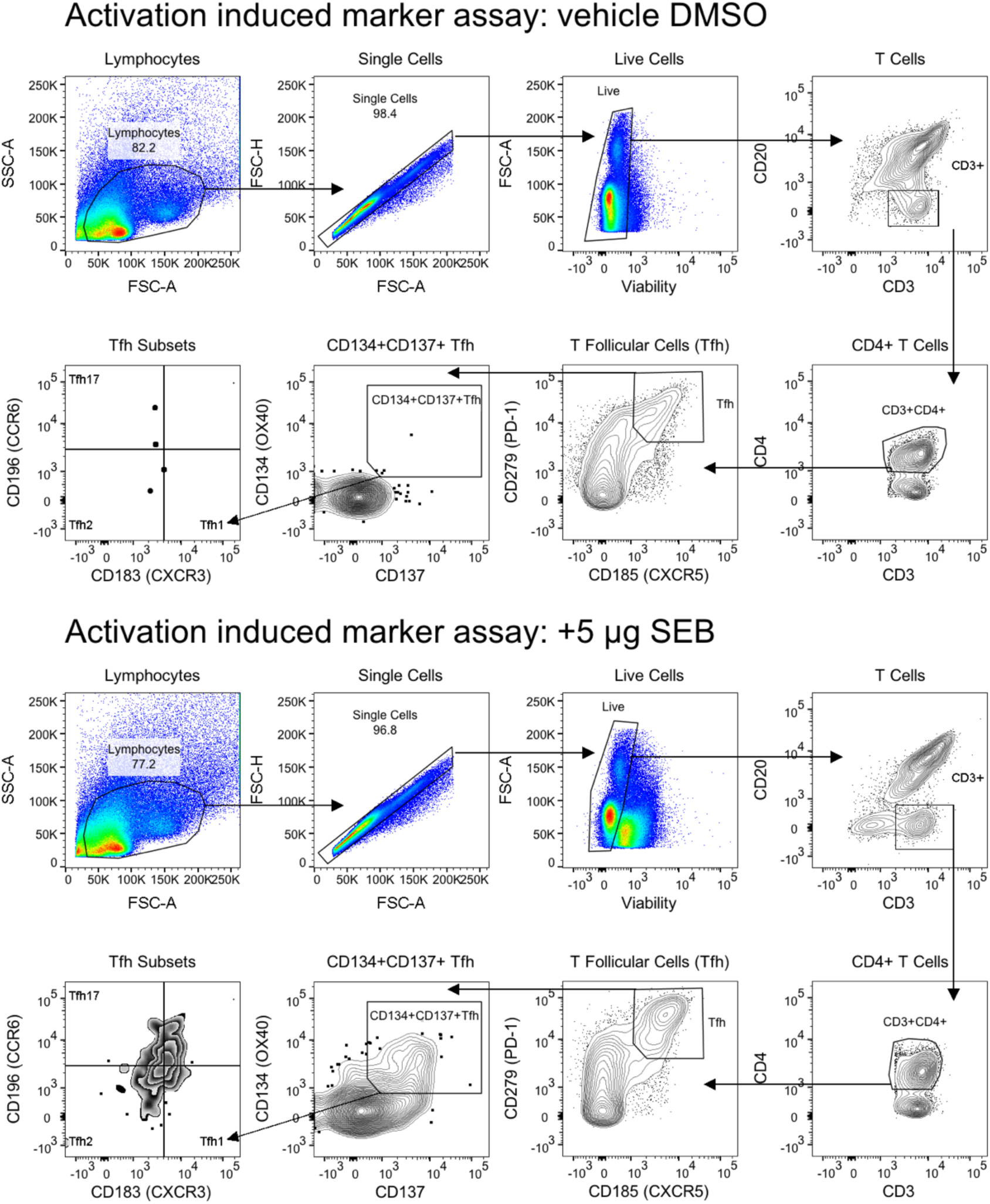
Activation-induced marker assay gating strategy for follicular T helper cells. Top two rows: DMSO vehicle control cultures; bottom two rows: staphylococcal enterotoxin B-stimulated samples.

### Supplementary tables

**Table S1.** Study Animals.

| Vaccine Group | Animal ID | Sex | Age at W0 (months) | Graphical Symbol |
| --- | --- | --- | --- | --- |
| Protein + 3M-052 | RM1 | Male | 2.6 | blue filled circle |
| Protein + 3M-052 | RM2 | Female | 2.4 | blue open circle |
| Protein + 3M-052 | RM3 | Male | 2.3 | blue filled triangle |
| Protein + 3M-052 | RM4 | Female | 2.3 | blue open triangle |
| Protein + 3M-052 | RM5 | Female | 2.2 | blue open inverted triangle |
| Protein + 3M-052 | RM6 | Male | 2.1 | blue filled inverted triangle |
| Protein + 3M-052 | RM7 | Female | 2.0 | blue open diamond |
| Protein + 3M-052 | RM8 | Male | 1.6 | blue filled diamond |
| mRNA-LNP | RM9 | Female | 2.5 | red open circle |
| mRNA-LNP | RM10 | Male | 2.4 | red filled circle |
| mRNA-LNP | RM11 | Male | 2.4 | red filled triangle |
| mRNA-LNP | RM12 | Female | 2.3 | red open triangle |
| mRNA-LNP | RM13 | Male | 2.2 | red filled inverted triangle |
| mRNA-LNP | RM14 | Female | 2.1 | red open inverted triangle |
| mRNA-LNP | RM15 | Male | 2.1 | red filled diamond |
| mRNA-LNP | RM16 | Female | 2.1 | red open diamond |

**Table S2.**
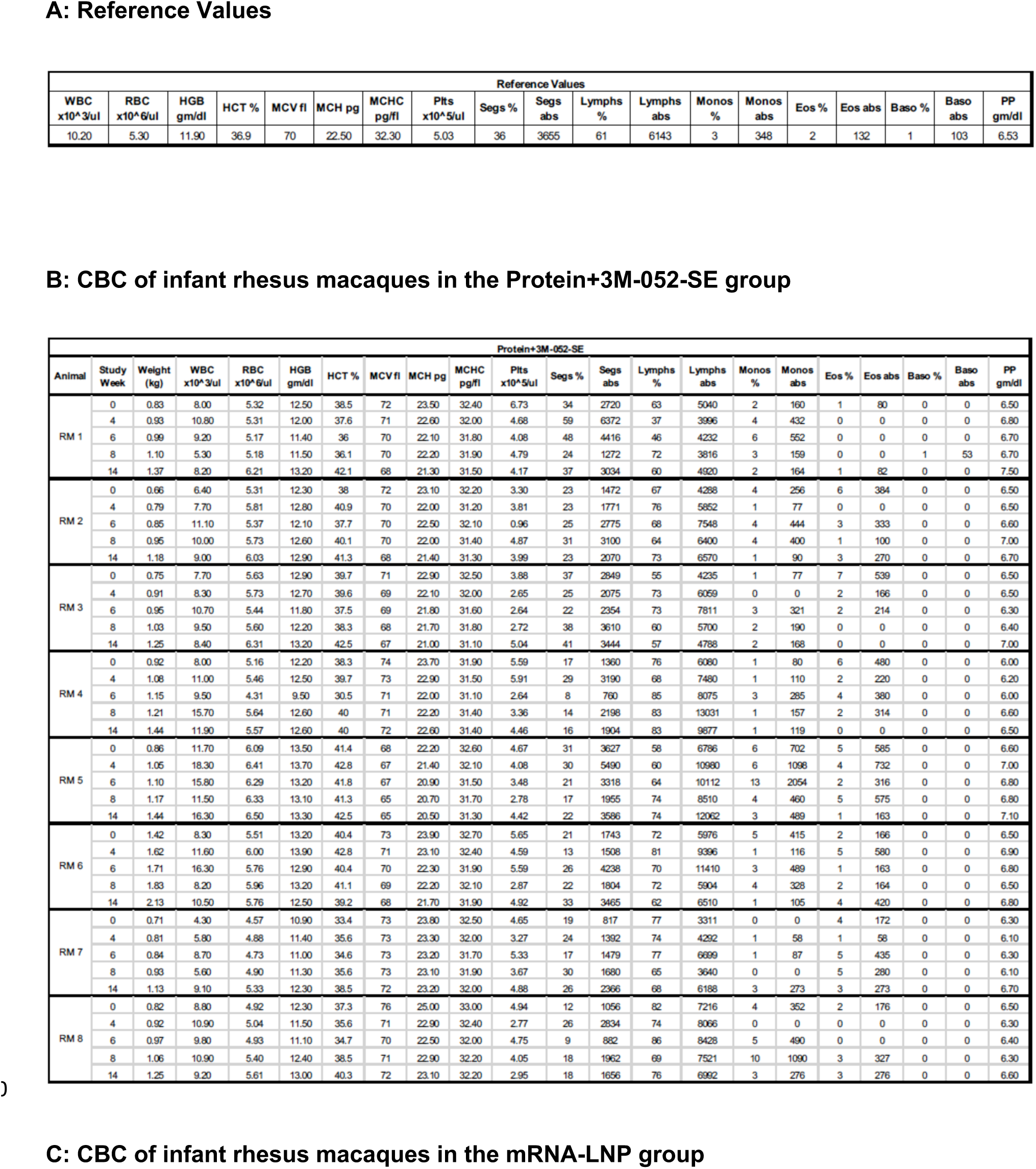
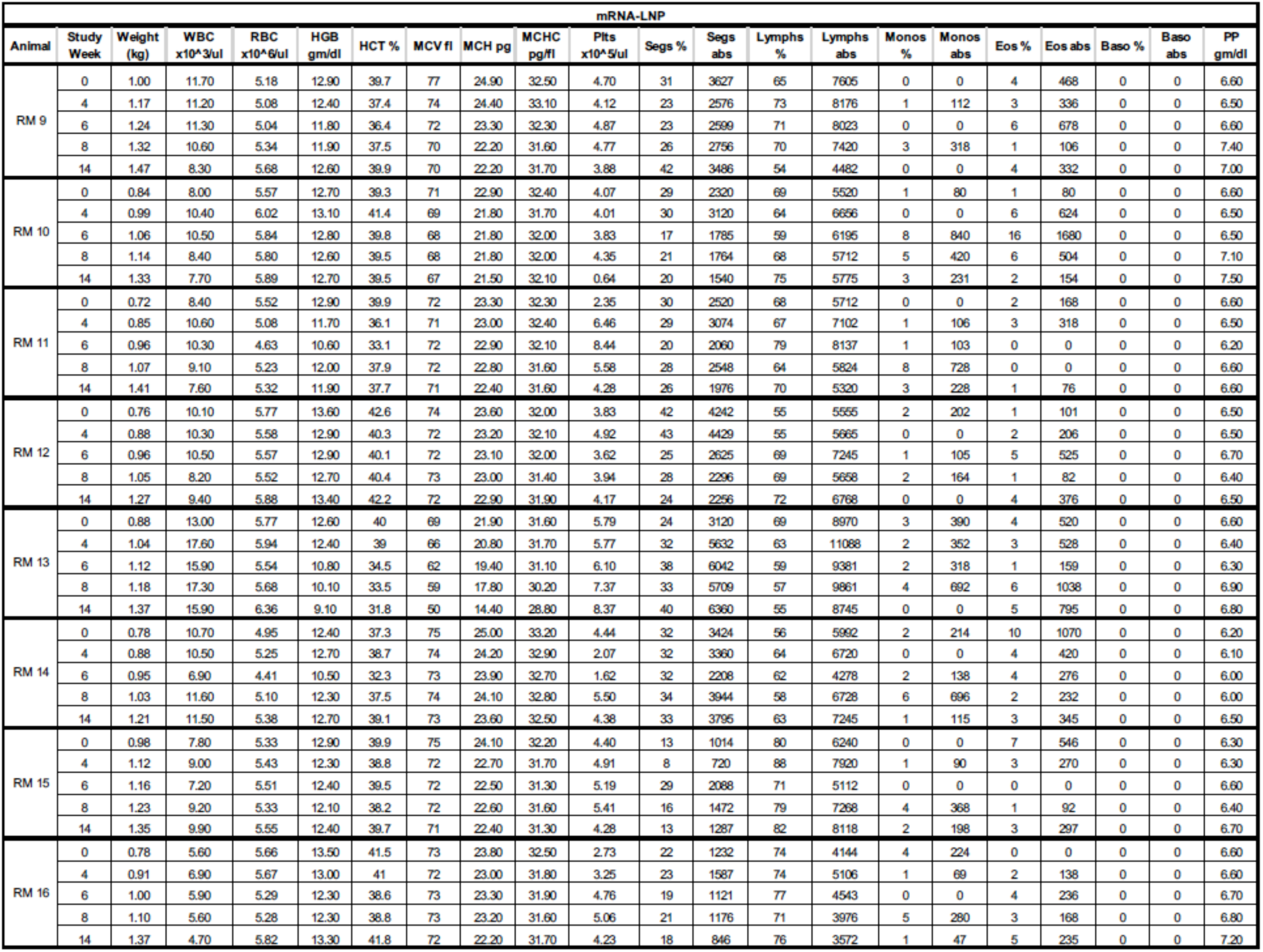
CBC reference values (A) for infant rhesus macaques age-matched to study animals that were immunized with the S-2P Protein+3M-052-SE (B) or the S-2P mRNA-LNP (C) vaccine.

**Table S3.**
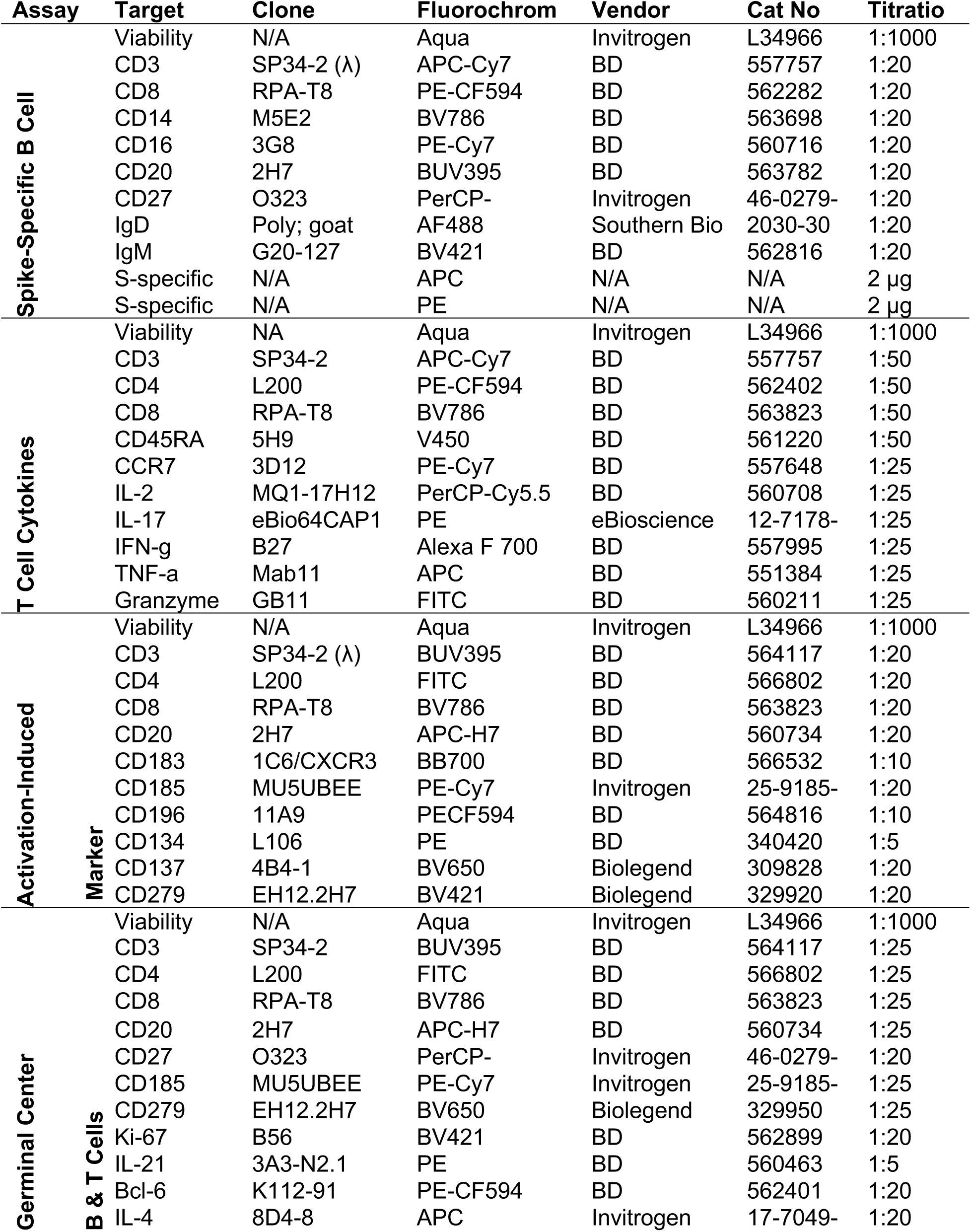
Reagents for Flow Cytometric Analysis.

## References

1. L. R. Baden et al., Efficacy and Safety of the mRNA-1273 SARS-CoV-2 Vaccine. N Engl J Med 384, 403–416 (2021).

2. A. T. Widge et al., Durability of Responses after SARS-CoV-2 mRNA-1273 Vaccination. N Engl J Med 384, 80–82 (2021).

3. E. E. Walsh et al., Safety and Immunogenicity of Two RNA-Based Covid-19 Vaccine Candidates. N Engl J Med 383, 2439–2450 (2020).

4. F. P. Polack et al., Safety and Efficacy of the BNT162b2 mRNA Covid-19 Vaccine. N Engl J Med 383, 2603–2615 (2020).

5. M. Voysey et al., Safety and efficacy of the ChAdOx1 nCoV-19 vaccine (AZD1222) against SARS-CoV-2: an interim analysis of four randomised controlled trials in Brazil, South Africa, and the UK. Lancet 397, 99–111 (2021).

6. J. Sadoff et al., Interim Results of a Phase 1-2a Trial of Ad26.COV2.S Covid-19 Vaccine. N Engl J Med, (2021).

7. C. Keech et al., Phase 1-2 Trial of a SARS-CoV-2 Recombinant Spike Protein Nanoparticle Vaccine. The New England journal of medicine 383, 2320–2332 (2020).

8. M. Guebre-Xabier et al., NVX-CoV2373 vaccine protects cynomolgus macaque upper and lower airways against SARS-CoV-2 challenge. Vaccine 38, 7892–7896 (2020).

9. Q. Bi et al., Epidemiology and transmission of COVID-19 in 391 cases and 1286 of their close contacts in Shenzhen, China: a retrospective cohort study. Lancet Infect Dis 20, 911–919 (2020).

10. A. H. Rowley, Understanding SARS-CoV-2-related multisystem inflammatory syndrome in children. Nature reviews. Immunology 20, 453–454 (2020).

11. L. A. Thompson, S. A. Rasmussen, One Year Later, How Does COVID-19 Affect Children? JAMA Pediatr 175, 216 (2021).

12. M. PrabhuDas et al., Challenges in infant immunity: implications for responses to infection and vaccines. Nature immunology 12, 189–194 (2011).

13. R. Rappuoli, Glycoconjugate vaccines: Principles and mechanisms. Sci Transl Med 10, (2018).

14. M. O. Ota et al., Hepatitis B immunisation induces higher antibody and memory Th2 responses in new-borns than in adults. Vaccine 22, 511–519 (2004).

15. D. K. Singh et al., Responses to acute infection with SARS-CoV-2 in the lungs of rhesus macaques, baboons and marmosets. Nature microbiology 6, 73–86 (2021).

16. K. S. Corbett et al., Evaluation of the mRNA-1273 Vaccine against SARS-CoV-2 in Nonhuman Primates. The New England journal of medicine 383, 1544–1555 (2020).

17. A. O. Hassan et al., A single intranasal dose of chimpanzee adenovirus-vectored vaccine protects against SARS-CoV-2 infection in rhesus macaques. bioRxiv, (2021).

18. K. Wu, et al., mRNA-1273 vaccine induces neutralizing antibodies against spike mutants from global SARS-CoV-2 variants. bioRxiv : the preprint server for biology, (2021).

19. S. P. Graham et al., Evaluation of the immunogenicity of prime-boost vaccination with the replication-deficient viral vectored COVID-19 vaccine candidate ChAdOx1 nCoV-19. NPJ vaccines 5, 69 (2020).

20. K. Abel, The rhesus macaque pediatric SIV infection model - a valuable tool in understanding infant HIV-1 pathogenesis and for designing pediatric HIV-1 prevention strategies. Curr HIV Res 7, 2–11 (2009).

21. G. Yiu et al., Evolution of ocular defects in infant macaques following in utero Zika virus infection. JCI Insight 5, (2020).

22. J. A. Mattison, K. L. Vaughan, An overview of nonhuman primates in aging research. Exp Gerontol 94, 41–45 (2017).

23. S. Reiss et al., Comparative analysis of activation induced marker (AIM) assays for sensitive identification of antigen-specific CD4 T cells. PloS one 12, e0186998 (2017).

24. A. B. Vogel et al., Immunogenic BNT162b vaccines protect rhesus macaques from SARS-CoV-2. Nature, (2021).

25. N. van Doremalen et al., ChAdOx1 nCoV-19 vaccine prevents SARS-CoV-2 pneumonia in rhesus macaques. Nature 586, 578–582 (2020).

26. D. J. Dowling et al., TLR7/8 adjuvant overcomes newborn hyporesponsiveness to pneumococcal conjugate vaccine at birth. JCI Insight 2, e91020 (2017).

27. B. Phillips et al., Adjuvant-Dependent Enhancement of HIV Env-Specific Antibody Responses in Infant Rhesus Macaques. J Virol 92, (2018).

28. B. Isho et al., Persistence of serum and saliva antibody responses to SARS-CoV-2 spike antigens in COVID-19 patients. Science immunology 5, (2020).

29. K. McMahan et al., Correlates of protection against SARS-CoV-2 in rhesus macaques. Nature, (2020).

30. I. Debock, V. Flamand, Unbalanced Neonatal CD4(+) T-Cell Immunity. Frontiers in immunology 5, 393 (2014).

31. M. E. Bottazzi, U. Strych, P. J. Hotez, D. B. Corry, Coronavirus vaccine-associated lung immunopathology-what is the significance? Microbes and infection 22, 403–404 (2020).

32. D. Deming et al., Vaccine efficacy in senescent mice challenged with recombinant SARS-CoV bearing epidemic and zoonotic spike variants. PLoS medicine 3, e525 (2006).

33. T. Bilich et al., T cell and antibody kinetics delineate SARS-CoV-2 peptides mediating long-term immune responses in COVID-19 convalescent individuals. Sci Transl Med, (2021).

34. T. Sekine et al., Robust T Cell Immunity in Convalescent Individuals with Asymptomatic or Mild COVID-19. Cell 183, 158–168.e114 (2020).

35. S. Belanger, S. Crotty, Dances with cytokines, featuring TFH cells, IL-21, IL-4 and B cells. Nat Immunol 17, 1135–1136 (2016).

36. Y. Cao et al., Potent Neutralizing Antibodies against SARS-CoV-2 Identified by High-Throughput Single-Cell Sequencing of Convalescent Patients’ B Cells. Cell 182, 73–84.e16 (2020).

37. G. E. Hartley et al., Rapid generation of durable B cell memory to SARS-CoV-2 spike and nucleocapsid proteins in COVID-19 and convalescence. Science immunology 5, (2020).

38. E. J. Anderson et al., Safety and Immunogenicity of SARS-CoV-2 mRNA-1273 Vaccine in Older Adults. The New England journal of medicine 383, 2427–2438 (2020).

39. A. D. Curtis, 2nd et al., A simultaneous oral and intramuscular prime/sublingual boost with a DNA/Modified Vaccinia Ankara viral vector-based vaccine induces simian immunodeficiency virus-specific systemic and mucosal immune responses in juvenile rhesus macaques. Journal of medical primatology 47, 288–297 (2018).

40. M. Dennis et al., Co-administration of CH31 broadly neutralizing antibody does not affect development of vaccine-induced anti-HIV-1 envelope antibody responses in infant Rhesus macaques. J Virol 93, e01783–01718 (2019).

41. L. Naldini et al., In vivo gene delivery and stable transduction of nondividing cells by a lentiviral vector. Science 272, 263–267 (1996).

42. Y. J. Hou et al., SARS-CoV-2 D614G Variant Exhibits Enhanced Replication ex vivo and Earlier Transmission in vivo. bioRxiv, (2020).

43. A. D. Curtis, 2nd et al., Oral Coadministration of an Intramuscular DNA/Modified Vaccinia Ankara Vaccine for Simian Immunodeficiency Virus Is Associated with Better Control of Infection in Orally Exposed Infant Macaques. AIDS research and human retroviruses 35, 310–325 (2019).

44. D. Finzi et al., Latent infection of CD4+ T cells provides a mechanism for lifelong persistence of HIV-1, even in patients on effective combination therapy. Nature medicine 5, 512–517 (1999).

45. J. D. Siliciano et al., Long-term follow-up studies confirm the stability of the latent reservoir for HIV-1 in resting CD4+ T cells. Nature medicine 9, 727–728 (2003).

46. M. J. Peluso et al., Differential decay of intact and defective proviral DNA in HIV-1-infected individuals on suppressive antiretroviral therapy. JCI insight 5, (2020).

47. D. Wrapp et al., Cryo-EM structure of the 2019-nCoV spike in the prefusion conformation. Science (New York, N.Y.) 367, 1260–1263 (2020).

48. P. A. Kozlowski et al., Modified wick method using Weck-Cel sponges for collection of human rectal secretions and analysis of mucosal HIV antibody. Journal of acquired immune deficiency syndromes (1999) 24, 297–309 (2000).

49. B. Phillips et al., Impact of Poxvirus Vector Priming, Protein Coadministration, and Vaccine Intervals on HIV gp120 Vaccine-Elicited Antibody Magnitude and Function in Infant Macaques. Clinical and vaccine immunology : CVI 24, (2017).

50. Y. Shaan Lakshmanappa et al., SARS-CoV-2 induces robust germinal center CD4 T follicular helper cell responses in rhesus macaques. Nature communications 12, 541 (2021).

51. L. Naldini, U. Blömer, F. H. Gage, D. Trono, I. M. Verma, Efficient transfer, integration, and sustained long-term expression of the transgene in adult rat brains injected with a lentiviral vector. Proceedings of the National Academy of Sciences of the United States of America 93, 11382–11388 (1996).

52. Y. J. Hou et al., SARS-CoV-2 D614G variant exhibits efficient replication ex vivo and transmission in vivo. Science (New York, N.Y.) 370, 1464–1468 (2020).

53. J. R. Whittle et al., Broadly neutralizing human antibody that recognizes the receptor-binding pocket of influenza virus hemagglutinin. Proceedings of the National Academy of Sciences of the United States of America 108, 14216–14221 (2011).

